# Deep exploration networks for rapid engineering of functional DNA sequences

**DOI:** 10.1101/864363

**Authors:** Johannes Linder, Nicholas Bogard, Alexander B. Rosenberg, Georg Seelig

## Abstract

Engineering gene sequences with defined functional properties is a major goal of synthetic biology. Deep neural network models, together with gradient ascent-style optimization, show promise for sequence generation. The generated sequences can however get stuck in local minima, have low diversity and their fitness depends heavily on initialization. Here, we develop deep exploration networks (DENs), a type of generative model tailor-made for searching a sequence space to minimize the cost of a neural network fitness predictor. By making the network compete with itself to control sequence diversity during training, we obtain generators capable of sampling hundreds of thousands of high-fitness sequences. We demonstrate the power of DENs in the context of engineering RNA isoforms, including polyadenylation and cell type-specific differential splicing. Using DENs, we engineered polyadenylation signals with more than 10-fold higher selection odds than the best gradient ascent-generated patterns and identified splice regulatory elements predicted to result in highly differential splicing between cell lines.

---

Designing DNA sequences for a target cellular function is a difficult task, as the cis-regulatory information encoded in any stretch of DNA can be very complex and affect numerous mechanisms, including transcriptional and translational efficiency, chromatin accessibility, splicing, 3’ end processing, and more. Yet, sequence-level design of genetic components and proteins has been making rapid progress in the past few years. Part of this advancement can be attributed to the collection of large biological data sets and improved bioinformatics modeling. In particular deep learning has emerged as state-of-the-art in predictive modeling for many sequence-function problems (Alipanahi et. al., 2015; Zhou et. al., 2015; Quang et. al., 2019; Avsec et. al., 2019; Kelley et. al., 2016; Greenside et. al., 2018; Kelley et. al., 2018; Jaganathan et. al., 2019; Cuperus et. al., 2017; Eraslan et. al., 2019). These models are now beginning to be combined with search heuristics and high-throughput assays to forward-engineer DNA and protein sequences (Rocklin et. al., 2017, Biswas et. al., 2018; Sample et. al., 2019; Bogard et. al., 2019). The ability to code regulatory DNA and protein function could prove useful for a wide range of applications. For example, controlling cell type-specific transcriptional, translational and isoform activity would enable engineering of highly specific delivery vectors and gene circuits. Functional protein design, e.g. generating heterodimer binders or proteins with optimally stable 3D structures, could prove transformative in T-cell therapy, drug interaction and drug delivery.

Discrete search heuristics such as genetic algorithms have long been considered the standard method for sequence design (Eiben & Smith, 2015; Shukla, Pandey & Mehrotra, 2015; Mirjalili et. al., 2020). Recently, however, gradient ascent optimization of the input sequence through a neural network fitness predictor has been proposed as a promising alternative. At its core, sequence generation via gradient ascent treats the input pattern as a position weight matrix (PWM). A neural network, pre-trained to predict a biological function, is used to evaluate the PWM fitness. The fitness score is used to compute a gradient with respect to the PWM parameters and the sequence PWM is iteratively optimized by gradient ascent. This class of algorithms, applied to sequences, was first employed to visualize transcription factor binding motifs learned from ChiP-Seq data (Lanchantin et. al., 2016). A modified version of the algorithm, with gradient estimators to allow passing sampled one-hot coded patterns as input, was used to engineer alternative polyadenylation (APA) sites (Bogard et. al., 2019). Direct gradient ascent on the input has also been successful in generating protein 3D structures (Evans et. al., 2018). Finally, the method has been used to indirectly optimize sequences with respect to a fitness predictor by traversing a pre-trained generative model and iteratively updating its latent input. For example, it has been applied to the input seed of a generative adversarial network (GAN) trained on the genome to engineer synthetic sequences that mimic conserved genomic elements (Killoran et. al., 2017).

Gradient ascent-style sequence optimization can be considered a continuous relaxation of discrete nucleotide-swapping searches, and as such makes efficient use of neural network differentiability; rather than naively trying out random changes, we follow a gradient to make stepwise local improvements on the fitness objective. Still, the basic method has a number of limitations. First, while the method makes incremental changes to all nucleotides simultaneously and may overcome some of the local minima a discrete search could not, it may nevertheless get stuck in local minima and the fitness of the converged patterns depend on PWM initialization (Bogard et. al., 2019). Second, it is computationally expensive to re-run gradient ascent for every sequence to generate. In fact, the method has no means of controlling the diversity of the optimized sequences, which may be required for generation of large candidate sequence sets.

To address these limitations, we developed Deep Exploration Networks (DENs), a variant of generative neural network models. The base architecture consists of a generator network connected to a differentiable fitness predictor. The generator produces sequence patterns which the predictor evaluates on the basis of an objective function; the overall goal is to generate sequences maximizing the objective. The core contribution of DENs is to explicitly control the sequence diversity generated during training. By making the generator compete with itself and penalize any two generated sequences based on similarity, we force the model to explore a much larger region of the cost landscape and effectively maximize both sequence fitness and diversity (**Figure 1A**). The architecture shares similarities with (Killoran et. al., 2017) but instead of optimizing the input seed of a pre-trained GAN, we optimize the weights of the generator to maximize both sequence fitness and diversity.

**Figure 1.**
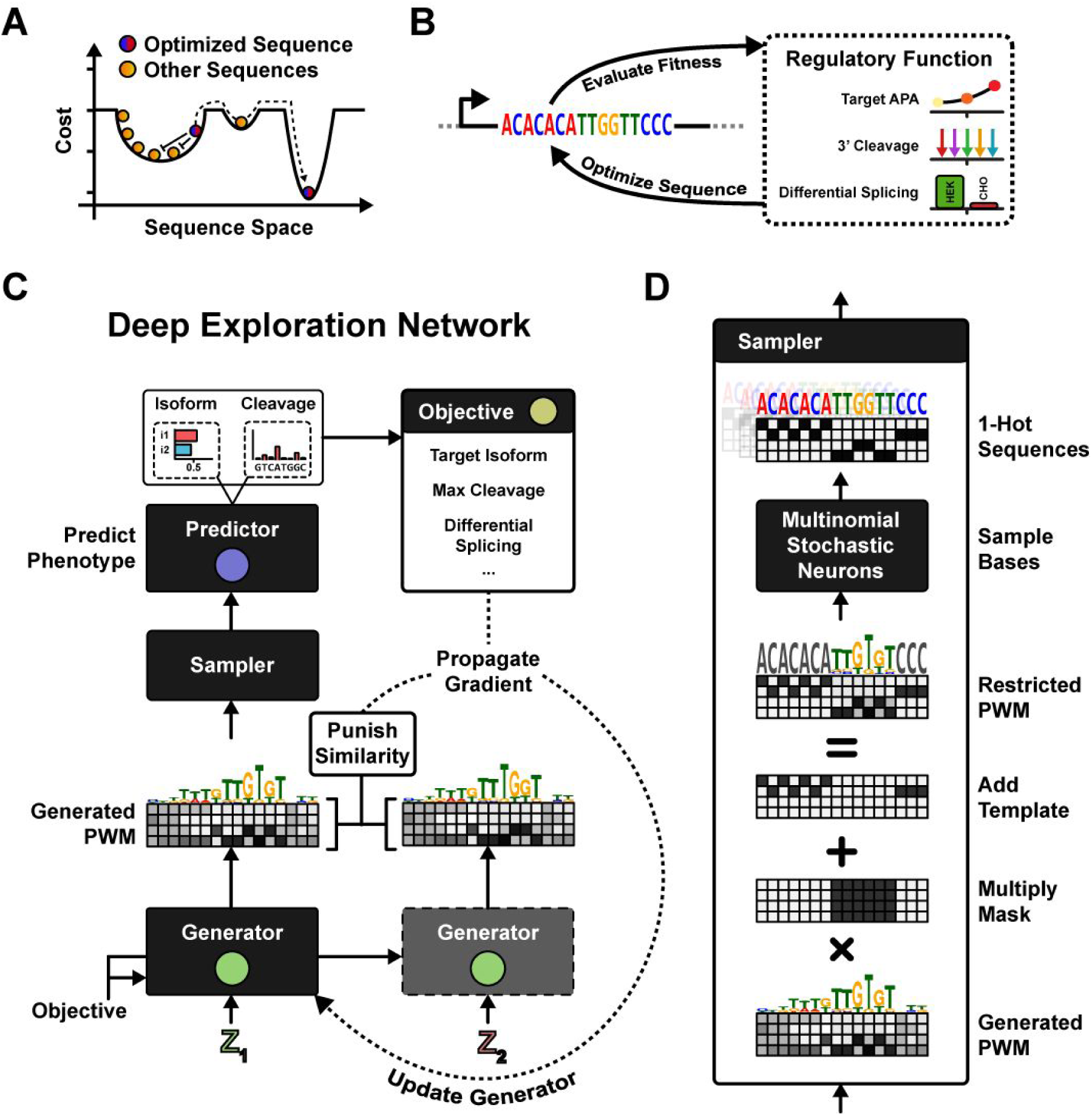
Deep Exploration Network Architecture. (A) A sequence produced by an input seed to a generative model (red/blue) shares the cost landscape with other generated sequences (orange). Patterns are penalized by similarity during training, resulting in an updated generator which transforms the red/blue seed into a different sequence, away from other patterns and potentially towards a new local minimum. (B) Sequences are optimized on the basis of a pre-trained fitness predictor to achieve some target function. This work focuses on three RNA isoform engineering applications: APA isoforms, 3’ cleavage positions, and differential splicing between two cell types. (C) In Deep Exploration Nets (DENs), the generator is run twice on two random seeds, producing two sequence PWMs. One of the PWMs is evaluated by the fitness predictor, resulting in an objective function gradient. The two PWMs are also punished by similarity, resulting in an exploration gradient, and the generator is updated by both gradients. (D) The PWM is multiplied by a mask (zeroing fixed nucleotides) and a template is added (encoding fixed letters). 1-hot-coded patterns are outputted by sampling nucleotides from stochastic neurons, and gradients are propagated by straight-through estimation.

Controlling pattern diversity enables DENs to sample multiple high-fitness outputs given a single input. To exemplify this idea, we construct “Inverse regression” models; given a real-valued regression target as input, the generator stochastically samples a sequence according to the target. This approach is conceptually similar to a variational autoencoder (Kingma & Welling, 2013) and conditioning by adaptive sampling (Brookes et. al., 2019), but rather than encoding the original pattern distribution or a conditional distribution, the model encodes the inverse of the predictor model while maximizing pattern variation.

We evaluate the utility of DENs on three synthetic biology applications (**Figure 1B**): First, we develop a basic model to generate 3’ UTR sequences with target APA isoform abundance. Second, we extend the model to do conditional multi-class generation in the context of guiding 3’ cleavage position. Finally, we use DENs to construct splice regulatory sequences that are predicted to result in maximal differential splicing between two cell types.

## Exploration In Deep Generative Models

The predictor 𝒫 used in a DEN is a differentiable model capable of predicting some property of an input pattern. The generator 𝒢 is a neural network designed to produce a pattern which can be passed as input to the predictor. Here, we are interested in generating DNA sequences; these patterns are typically represented as 1-hot-coded matrices, where the columns (length N) denote nucleotide position and rows (M channels) denote nucleotide identity. Hence, the pattern space is {0, 1}^*N*×*M*^. The predictor output is used to define an objective (the *cost function*), and the overall goal is to optimize the generator such that the generated sequences minimize the cost function (**Figure 1C**). The predictor thus provides gradients of the objective to the generator, which in turn can update its internal weights by gradient descent. At convergence, the generator is optimized to synthesize new patterns which are (locally) optimally minimizing the cost. Only the generator is optimized, having pre-trained the predictor network to accurately predict the targeted biological function.

We first define the fitness objective in terms of the predictor’s output. For example, in the case of models that predict isoform abundances, we might want to minimize the KL-divergence between the predicted and target isoform proportion. However, if we only maximize the fitness objective, it is likely that the generator will learn to only produce one single pattern, even when feeding new input seeds to the generator. The generator can simply choose to ignore the entropy induced by the seed. In fact, if the cost function has many local minima, there is nothing promoting exploration during training of the generator. Rather, it learns to always output the pattern located at the bottom of the local optimum we started in. There may however exist other, better, local minima.

The distinguishing feature of a DEN is to enforce exploration during training by controlling the degree of sequence diversity generated by the network. Diversity is explicitly controlled in the cost function by making the generator compete with itself; we penalize any two generated sequence patterns based on similarity, forcing the generator to maintain entropy in the weights from the input seed and constantly form new sequences. This mechanism is implemented by running the generator twice at each step of the optimization, with two random seeds, and penalizing the two patterns on the basis of a similarity metric. We refer to the coefficient of the similarity loss in the cost function as the *repel weight*.

This cost function layout is quite different compared to a classical GAN (Goodfellow et. al., 2014), which is typically optimized to minimize some cost C(𝒟(Data), 𝒟(𝒢(*z*))) such that an adversarial discriminator 𝒟 can not distinguish between the real data and the distribution generated by 𝒢. Here, we instead jointly minimize the fitness cost C_fitness_(𝒫(𝒢(*z*_1_)) of an arbitrary predictor 𝒫 and an adversarial diversity cost C_diversity_(𝒢(*z*_1_), 𝒢(*z*_2_)) of the generator 𝒢. Note that, in contrast to (Killoran et. al., 2017) where optimization is done on a single input seed of a pre-trained GAN, min_*z*_ C_fitness_ (𝒫(𝒢(*z*))), we optimize the generator 𝒢 itself, min_𝒢_ C_fitness_ (𝒫(𝒢(*z*))) + C_diversity_(𝒢(*z*_1_), 𝒢(*z*_2_)) for all seeds *z*_1_, *z*_2_ ∈ *U*(0, 1)^100^.

In the context of genomics, where the generator produces 1-hot-coded sequences, we penalize patterns using a multi-offset cosine similarity metric. We found empirically that minimizing a slack-bound cosine similarity gives the best results, where a fraction of the sequences can be identical up to a margin *ϵ* without incurring any loss. Given two patterns *X*^(1)^ = 𝒢(*z*^(1)^) and *X*^(2)^ = 𝒢(*z*^(2)^) generated by 𝒢, we define the similarity loss as:

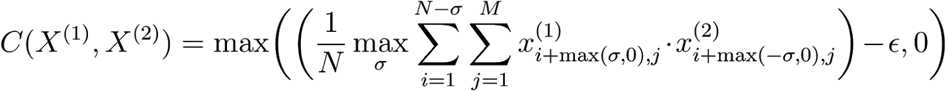

The cost function now provides gradients rewarding diverse pattern generation and this exploration component is balanced by the exploitation component of maximizing the fitness objective. The generator can be trained to minimize the compound cost using a gradient-based optimizer. We built the DEN in Keras (Chollet et. al., 2015) and optimized the generator with Adam (Kingma et. al., 2014).

## Pattern Representation for Genomics

A generator cannot output discrete 1-hot-coded patterns in {0, 1}^*N*×*M*^ and still maintain differentiability from the predictor to the generator. Two different methods have been proposed to address this issue: (1) representing the input pattern as a continuous, differentiable distribution, and (2) representing the pattern by discrete samples and approximating the gradient. Both methods are coupled with their own intrinsic artifacts and we show that using both representations together during training may enhance convergence.

In both methods, the generator produces patterns in ℛ^*N*×*M*^, representing nucleotide scores. By applying a column-wise Softmax transform, we turn these scores into nucleotide probabilities (a PWM). Finally, we multiply the PWM with a mask matrix, zeroing out all elements corresponding to fixed (non-changeable) sequence context, and add a template matrix which encodes the fixed sequence (**Figure 1D**). The first method has previously been demonstrated in the context of genomics (Killoran et. al., 2017; Stewart et. al. 2018) and we refer to it here as Relaxed Input Form Representation (RIFR). In RIFR, the PWM is directly passed to the predictor (as a continuous relaxation [0, 1]^*N*×*M*^ of the input). The predictor has never been trained on real-valued patterns and may perform poorly on high-entropy PWMs. However, we can push the PWMs toward a 1-hot-coded state during training by minimizing PWM entropy in the cost function. Empirically, we found that minimizing an absolute error between the average nucleotide entropy and a target entropy works well. Note that minimizing PWM entropy does not mean that we necessarily minimize generator entropy. On the contrary, if we promote a high degree of exploration by punishing similar PWMs, the generator learns to produce diverse, low-entropy PWMs.

RIFR has a fundamental drawback: The gradient propagated backward through the PWM quickly approaches zero as the nucleotide logits push the Softmax probabilities toward their extremes. The problem is exacerbated as we explicitly minimize PWM entropy. Put differently, we optimize the system for vanishing gradients, resulting in halted convergence.

In the second method, which we here call Sampled Input Form Representation (SIFR), we circumvent the Softmax representation by taking advantage of sampling techniques and straight-through (ST) gradient estimators (**Figure 1D**; Bengio, Léonard & Courville, 2013; Courbariaux et. al., 2016; Bogard et. al., 2019). We use the generated nucleotide logits as parameters of a multinomial probability distribution, from which we draw K independently sampled 1-hot-coded patterns. The K samples are used as input to the predictor and the average gradient is propagated backwards through the sampling distribution using ST estimation. While the increased sample variance is an obvious drawback, it can be mitigated by increasing the number of samples drawn at each step. However, optimization can be noisy even with infinitely many samples (K→inf), since gradients produced by ST estimation may at times be incorrectly approximated. As our results indicate below, combining both methods and walking down the average gradient (Dual Input Form Representation, or DIFR) can reduce variance and estimation artifacts while overcoming vanishing gradients.

## Engineering APA Isoforms

We first demonstrate deep exploration nets in the context of Alternative Polyadenylation (APA). APA is a post-transcriptional 3’ end processing event where competing polyA signals (PAS) in the same 3’ UTR give rise to multiple mRNA isoforms (**Figure 2A**) (Di Giammartino et al., 2011; Tian and Manley, 2017). A typical PAS consists of a core sequence element (CSE), often the hexamer AATAAA, as well as diverse upstream and downstream sequence elements (USE, DSE). Cleavage and polyadenylation occurs approximately 17 nt downstream of the CSE within the DSE. In a competitive situation with multiple PASs in the same 3’UTR, the sequence of each PAS is the major determinant of isoform selection.

**Figure 2.**
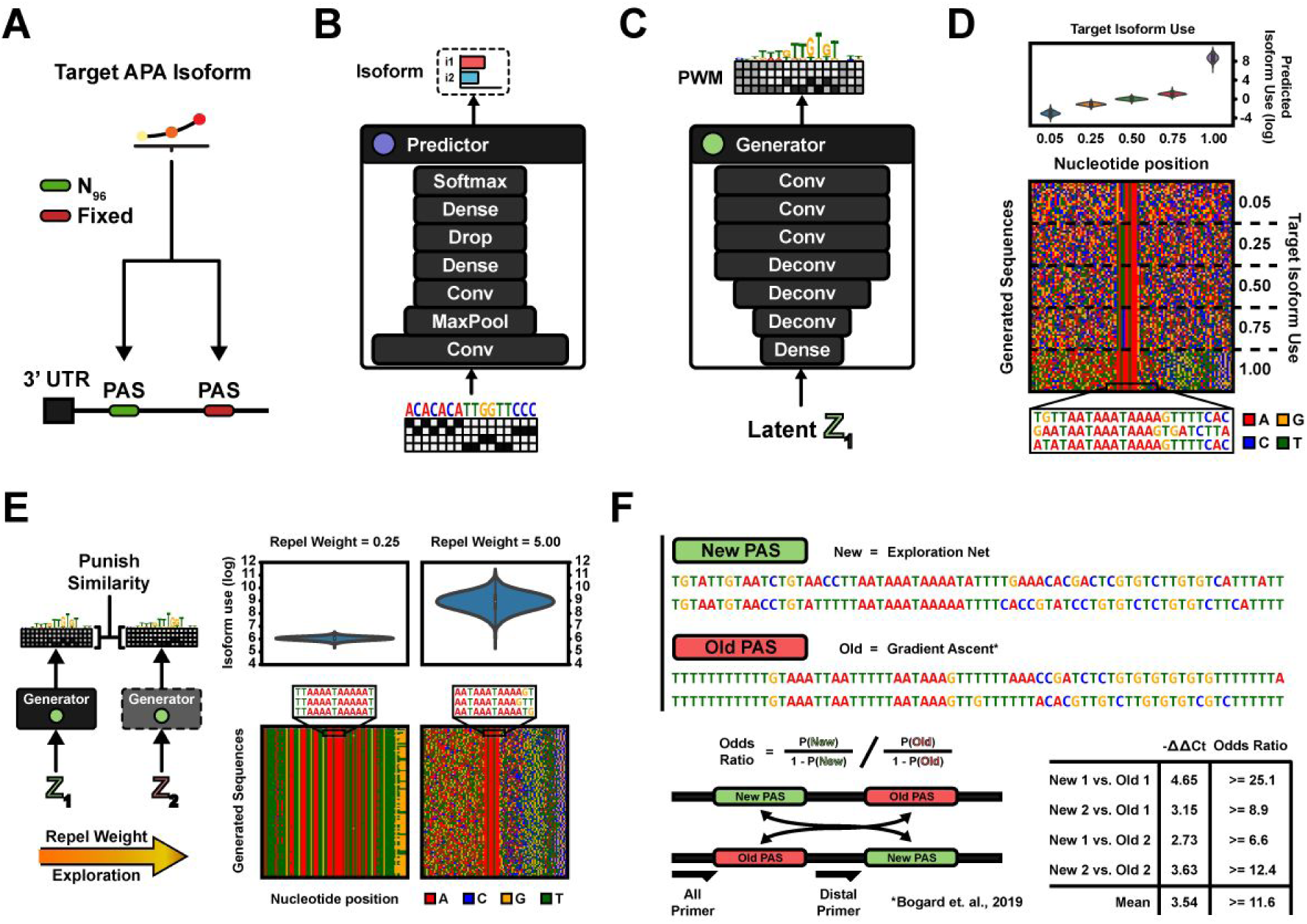
Engineering APA Isoforms. (A) Two PASs in a 3’ UTR compete for cleavage and polyadenylation. The generative task is to design proximal PASs which are selected at a target proportion. (B) The APA predictor architecture. A set of convolutional, pooling, dropout and dense layers transform the 1-hot-coded input sequence into an APA isoform proportion prediction. (C) The generator follows a typical GAN architecture. Dense and (de-)convolutional layers transform the input seed vector into a sequence PWM of nucleotide log probabilities. (D) Evaluation of five separate DENs trained to generate sequences according to APA isoform targets: 5%, 25%, 50%, 75% and 100% (‘Max’). (Top) Predicted isoform proportions of 1,000 sampled sequences per target objective. Mean and Std dev of isoform log odds per target (proportions in parenthesis): (Target 5%) −2.99 +- 0.46 (= 5.25% +- 2.42%), (Target 25%) −1.12 +- 0.30 (= 25.06% +- 5.66%), (Target 50%) 0.026 +- 0.26 (= 50.6% +- 6.3%), (Target 75%) 1.08 +- 0.31 (74.2% +- 5.79%), (Target Max) 8.68 +- 0.72 (99.98% +- 0.02%). (Bottom) Generator sequence diversity, illustrated by 20 randomly sampled sequences per objective on a pixel grid where rows denote sequences and columns nucleotide position. 0% duplication rate at 100,000 sampled sequences by any of the generators. Hexamer entropy ranges between 9.11 and 10.0 bits depending on generator (of 12 bits maximum), with 2,134 to 3,203 unique hexamers across the first 1,000 sequences depending on generator. (E) The sequence similarity loss was evaluated by re-training the Max-target APA isoform generator, in one instance with a low similarity loss coefficient (left) and in another instance a high coefficient (right). (Top) Predicted isoform proportions for 1,000 generated sequences, per generator instance. (Bottom) Sequence diversity illustrated by sampling 100 sequences. Low repel weight (left): Mean predicted isoform log odds = 6.06 +- 0.12, and 99.5% duplication rate at 100,000 sampled sequences. High repel weight (right): Mean predicted isoform log odds = 8.91 +- 0.72, and 0% duplication rate at 100,000 samples. (F) Experiment validating the performance of the generated sequences. Two Max-target sequences generated by the DEN were synthesized as either the proximal or distal pA signals on a minigene reporter in competition with baseline gradient ascent-generated sequences using the same fitness predictor (Bogard et. al., 2019). Isoform odds ratios (preference fold changes) were assayed using qPCR and estimated from cycle threshold values. The newly generated sequences have on average 9.4-fold increased preference.

We previously developed a neural network for predicting APA isoform abundance (APARENT; Bogard et. al., 2019), which we use here as the predictor (**Figure 2B**). The DEN was tasked with generating PASs with precisely defined target isoform abundances as well as maximally strong PASs. We have previously generated such sequences using direct gradient ascent and experimentally validated them (Bogard et. al., 2019), enabling benchmark comparisons. The generator chosen for this application follows a DC-GAN architecture (Radford et. al., 2015; **Figure 2C**). When training the generator, we pass both the PWM and a number of sampled one-hot patterns as input to the predictor, walking down the average loss gradient (DIFR).

We trained 5 instances of the generator, each optimized to generate sequences according to the following target isoform proportions: 0%, 25%, 50%, 75% and maximal use (‘Max’). These objectives were encoded in the cost function by minimizing the KL-divergence between the predicted APA isoform proportion and the target proportion (F**igure S2A**). After training, each generator could produce sequence samples fulfilling its target isoform proportion with high precision (**Figure 2D** Top): Each generated isoform distribution mean was within 1% from the target proportion, with a maximum standard deviation of 5.79%. The generated sequences for the Max-objective were predicted to be extremely efficient PASs (on average 99.98% predicted use with less than 0.02% deviation). All five generators exhibited a high degree of diversity (**Figure 2D** Bottom, **S2B-C**); when sampling 100,000 sequences per generator, no two sequences were ever identical (0% duplication rate) and all generators had a hexamer entropy of between 9.11 and 10.0 bits (of 12 bits maximum), with up to 3,203 unique hexamers in the first 1,000 sampled sequences. We replicated the entire analysis for polyA signals with a different 3’ UTR context (**Figure S2D**), showing that the method can easily be adapted to new contexts by re-configuring the generator; we simply changed the generator mask to zero out positions where the sequence is fixed, and changed the template to encode the new fixed sequence (**Figure 1D**).

To evaluate the importance of promoting exploration while training, we re-trained the Max isoform-generator with two different parameter settings; in one training instance, we lowered the repel weight (the similarity loss coefficient) to a small value, and in another instance we increased the repel weight (**Figure 2E, S2E**). With a low repel weight, the generator only learns to sample few, low-diversity sequences, all of similar isoform log odds (**Figure 2E** Left; mean isoform log odds = 6.06, 99.5% duplication rate at 100,000 samples). With an increased repel weight, generated sequences become much more diverse and the mean isoform odds increase almost 20-fold (**Figure 2E** Right; mean isoform log odds = 8.91, 0% duplication rate at 100,000 samples). These results indicate that exploration during training drastically improves the final fitness of the generator. We further evaluated the Max-isoform generator when using one-hot samples (SIFR), the continuous PWM (RIFR), or a combination of both (DIFR) as input to the predictor (**Figure S2F**). Using DIFR, the loss significantly improves, with less than 50% the magnitude of the RIFR loss after 30 epochs. Finally, we validated the accuracy of the target-isoform generators against the MPRA datasets published in (Bogard et. al., 2019), by comparing the generated sequences against sequences with known isoform ratios estimated from RNA-Seq (**Figure S2G-H**).

## Experimental Validation of Deep Exploration-Sequences

As suggested in **Figure 2E**, exploration increases the capability of generating high-fitness sequences. Next, we wanted to characterize experimentally whether DEN-generated PASs truly are stronger (more optimal) than sequences generated by the baseline gradient ascent method and, if so, how much stronger they are. To that end, we synthesized APA reporters with two adjacent PASs (**Figure 2F**): Each reporter contained one of the newly generated Max-target PASs, as well as one of the strongest gradient ascent-optimized signals from (Bogard et. al., 2019). In order to discount first-come-first-serve bias, we experimentally assayed both signal orientations for each reporter. The reporters were cloned onto plasmids and delivered to HEK293 cells. We quantified the expressed RNA isoform levels using a qPCR assay, measuring the Ct values of total and distal RNA respectively. Using Ct differences to estimate odds ratio lower bounds, we found that the DEN-generated sequences were on average 11.6-fold more preferred (usage odds increase) than the gradient ascent-generated sequences (**Figure 2F**). To put this in perspective, the strongest gradient ascent-sequence had usage odds of 127:1 (99.22%) relative to a distal bGH PAS separated by 200 nt. The DEN-sequences would have usage odds of 1481:1 (99.93%) relative to the same signal.

## An Inverse Regression Model of APA

Isoform prediction is a continuous regression problem (predicting proportions in the continuous range from 0 to 1), yet in the above modeling framework we discretize and hardcode one specific isoform proportion to target per generator network. However, exploration networks allow us to capture the entire distribution of isoform proportions by adding a “Target isoform proportion”-input, effectively turning the generator into an inverse regression model capable of stochastically sampling a sequence given a target regression value (**Figure 3A, S3A-B**). During training, we randomly sample target isoform proportions, feed it to the generator as input and simultaneously specify the same target isoform proportions in the loss function (which we compare the predictor model to). As a result, we train the generator to sample sequences which fulfill whatever target isoform proportion was supplied as input. Our results show that this training scheme works remarkably well (**Figure 3B**); when sampling 10,000 sequences from the generator, the generated sequences’ predicted isoform log odds were highly correlated with their corresponding targets (pearson r = .97, p = 0), and had a sequence duplication rate of 0%, meaning we could successfully generate diverse sequences which, according to the predictor, satisfied the inverse regression objective. When projecting the sequences in two-dimensional space using tSNE (Maaten et. al., 2008), we find only a single cluster, where the sequences smoothly transition from one edge to the other based on the isoform log odds.

**Figure 3.**
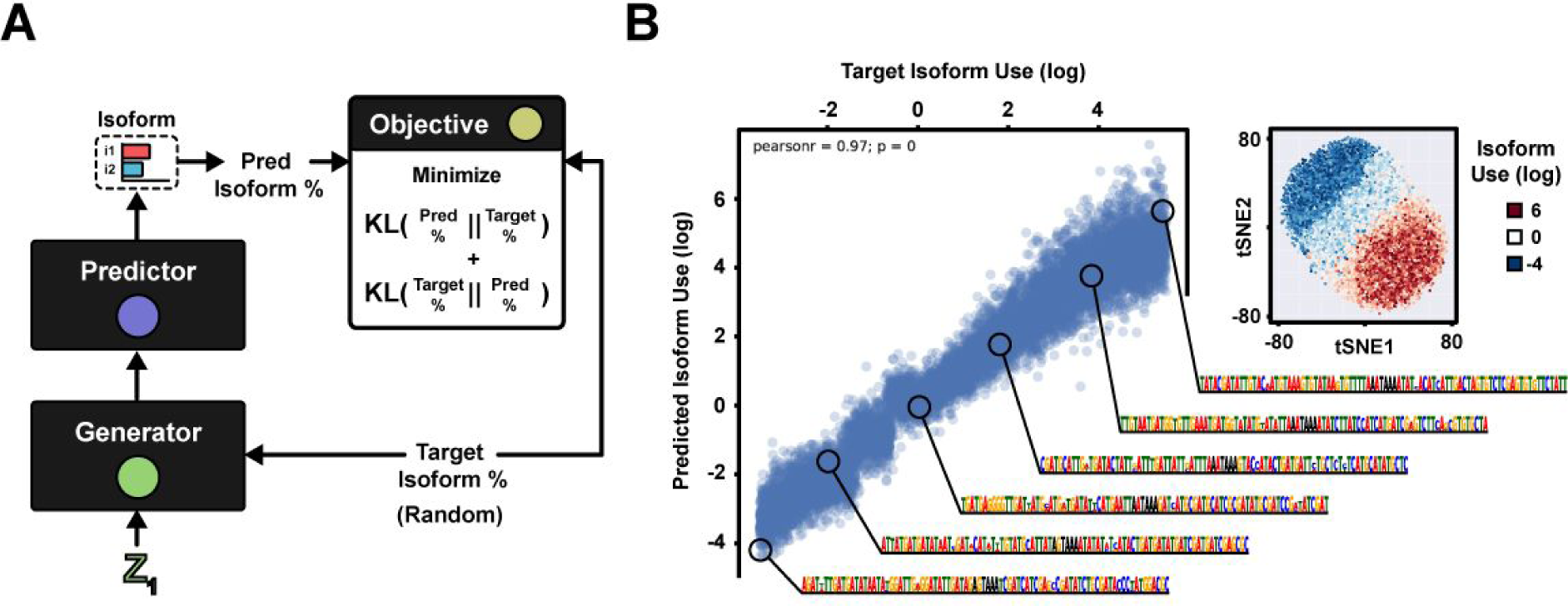
An Inverse Regression Model. (A) The DEN is trained by randomly sampling target isoform proportions and passing them to both the generator and objective. The generator is optimized to produce sequences which are predicted to conform to the sampled target proportions. (B) A deep exploration network, using the generator architecture of Figure 3A, is trained to generate PASs according to randomly sampled target isoform logits in the range −4 to 6. After training, the predicted isoform logits of the generated sequences strongly agree with their corresponding target logits (pearson r = 0.97, n = 10,000). The generated sequences are clustered in tSNE and colored according to their target isoform logits.

## Engineering 3’ Cleavage Position

The next application is closely related to APA, but rather than multiple competing PASs, we here concern ourselves with the position of 3’ cleavage within a single signal (**Figure 4A**). Cleavage occurs downstream of the central polyadenylation element – the CSE hexamer – however the exact position and magnitude is tightly regulated by a complex code (Elkon et al., 2013). The predictor model used for APA above – APARENT – can also predict the 3’ cleavage distribution and so we re-use the model here (**Figure S4A-B**). Tasked with generating sequences which maximize cleavage at 9 distinct positions, we constructed a multi-class exploration network with a generator architecture similar to class-conditional GANs (Mirza et al., 2014; **Figure 4B, S4C**), where an embedding layer transforms the class label (target cut position) into a high-dimensional vector which is concatenated onto every layer of the generator. The loss remains the same as before, except now the predicted cleavage distribution is used to minimize KL-divergence against the target distribution.

**Figure 4.**
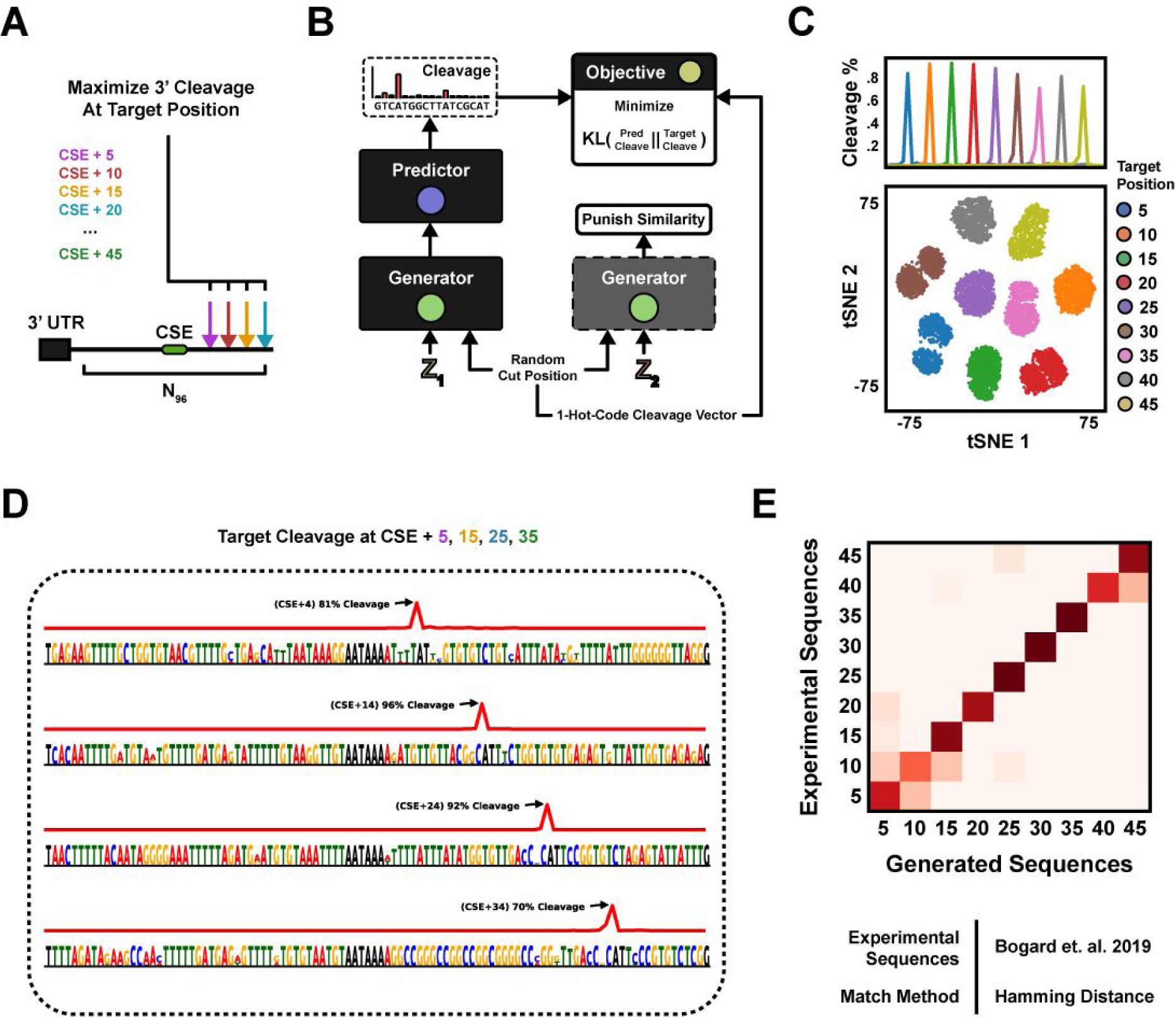
Engineering 3’ Cleavage Positions. (A) The 3’ mRNA cleavage position is governed by a cis-regulatory code within the PAS. The generative task is formulated as designing PASs which maximize cleavage at target nucleotide positions downstream of the central hexamer (CSE) AATAAA. (B) The DEN is trained by randomly sampling cut positions along with the seeds, and passing the cut positions to both the generator and objective. The generator is optimized to produce sequences which are predicted to cleave according to the target cut positions. (C) Evaluation of a class-conditional DEN trained to generate sequences with maximal cleavage at 9 positions, +5 to +45nt downstream of the CSE. (Top) Mean predicted cleavage profile of 1,000 sampled sequences per target position. X-axis denotes nt position and Y-axis is cleavage %. Predicted vs. target cut position R^2 = 0.998. (Bottom) All 9,000 sequences were clustered in tSNE and colored according to their target cleavage position. (D) Example sequences generated by the network for target positions +5, +15, +25 and +35. Sequence generation is highly diverse, with 0% duplication rate at 100,000 samples. Hexamer entropy = 9.07 of 12 bits, with 1,727 unique hexamers across 1,000 sequences. (E) The newly generated sequences were compared against gradient ascent-generated sequences for the same target (Bogard et. al., 2019), by defining each cluster centroid as the mean one-hot pattern of all DEN-generated sequences and assigning each gradient ascent-pattern to the closest centroid based on L1 distance. Agreement = 0.87.

After training, the generator could sample diverse sequences with highly specific cleavage distributions given an input target position (**Figure 4C-D**; Predicted vs. target cut position R^2 = 0.998, 0% duplication rate at 100,000 sampled sequences; S4D-F, Movie S1-S2). When clustering the sequences in tSNE (**Figure 4C**, bottom), we observe clearly separated clusters based on the target cleavage position. We further confirmed the function of the sequences by comparing them to a set of gradient ascent-optimized sequences which had previously been validated experimentally with RNA-Seq (**Figure 4E**; Nearest Neighbor-agreement = 87%) (Bogard et. al., 2019).

## Engineering Cell type-specific Differential Splicing

While precise cis-regulatory control in a single cell type has important applications, one of the hardest yet perhaps most interesting problems in genomics and synthetic biology is to code cis-regulatory functions which are differentially expressed across multiple cell types. Here we consider the task of engineering cell type-specific differential splice forms (**Figure 5A**; Blencowe, 2006; Roca, Krainer & Eperon, 2013; Lee & Rio, 2015). Specifically, we define the task as maximizing the difference in splice donor usage for an alternative 5’ splicing event in two different cell lines, by designing the regulatory sequences (25nt) downstream of each alternative donor. This particular splicing construct has been studied in the context of HEK293 cells (Rosenberg et. al., 2015), where MPRA data measuring hundreds of thousands of variants were collected. To study differential effects across cell types, we report new MPRA measurements of this splicing library in HELA, MCF7 and CHO cells, which we used together with the original HEK data to train a cell type-specific 5’ splice site usage prediction network (**Figure 5B, S5A**). The trained network could accurately predict splicing isoform proportions on a held-out test (**Figure S5B**; mean R^2 = 0.88). Importantly, the predicted difference in splice site usage between cell types had a strong correlation with measured differences (**Figure S5C**; predicted vs. measured dPSI R^2 ranged between 0.35 and 0.47 depending on cell type pair). We focused on MCF7 and CHO, as the largest average differential trend was observed between these two cell lines.

**Figure 5.**
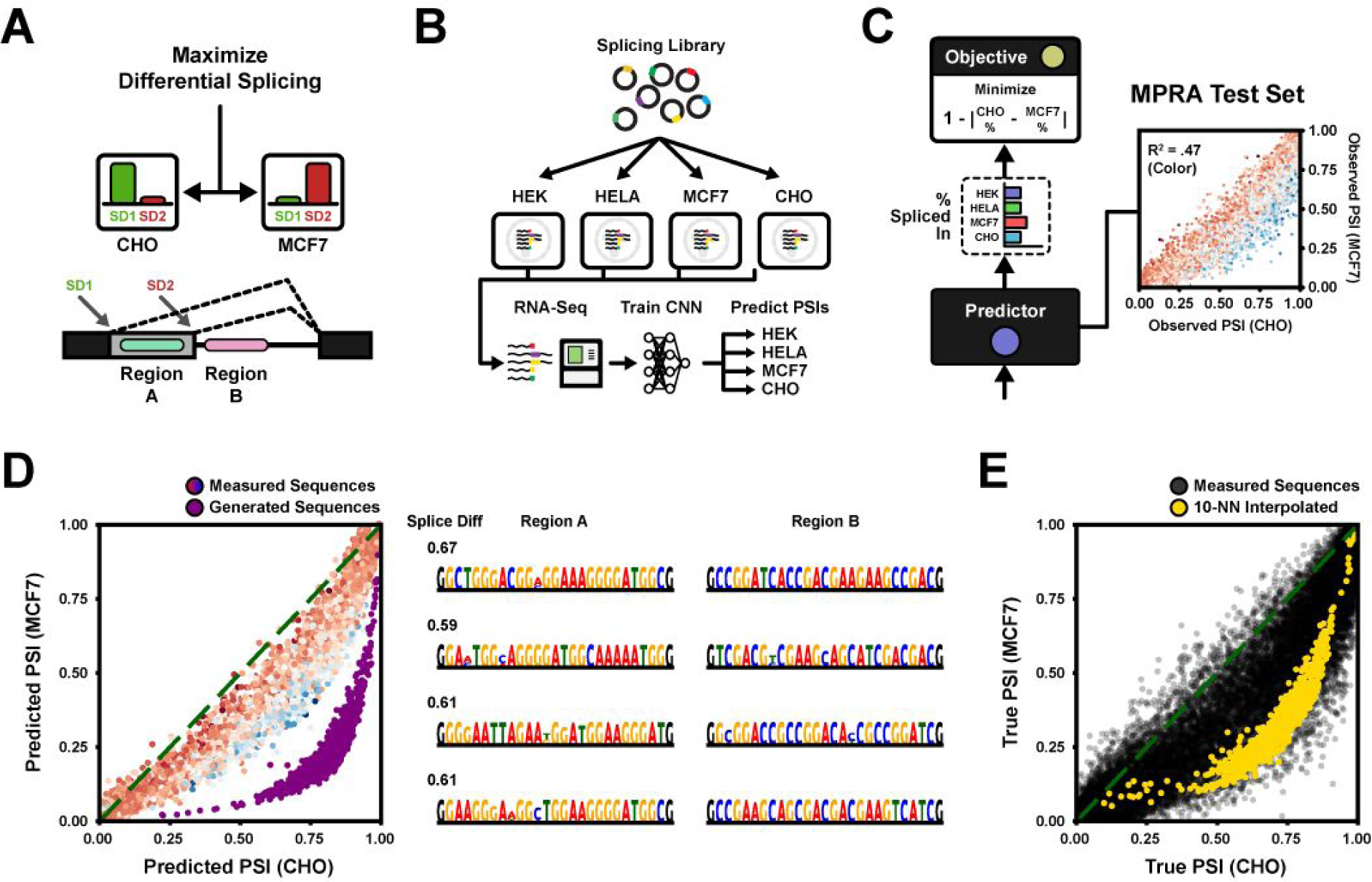
Engineering Differential Splicing Across Cell Types. (A) Two 5’ splice donors compete for splicing. The task is to design two sequence regions, Region A which is located between the donors and Region B which is located downstream of the 3’-most donor, to maximize differential usage (PSI) of donor 1 between two cell types. (B) Summary of the experimental pipeline. The MPRA of (Rosenberg et. al., 2015) was originally measured in HEK293 cells. Here, the library was transfected in additional cell lines HELA, MCF7 and CHO and measured by RNA-Seq. A neural network (CNN) was trained on all four cell line datasets to predict PSI per cell type given only the DNA sequence as input. (C) Left: The predicted MCF7 and CHO PSIs are used to maximize absolute difference. Right: Measured MPRA test set PSIs for MCF7 and CHO. Color indicates predicted dPSI (blue/red = more/less used in CHO). Predicted vs. measured dPSI R^2 = 0.47. (D) DEN evaluation on the basis of the prediction network. (Left) Predicted PSIs in MCF7 and CHO for 1,000 sequences sampled from the trained generator, plotted together with predicted MPRA test set PSIs in MCF7 and CHO. Color indicates measured dPSI of test sequences. Purple dots indicate generated sequences. Mean predicted dPSI of test sequences = 0.08 (+- 0.07). Mean predicted dPSI of generated sequences = 0.56 (+- 0.07). (Right) Selection of maximally differentially spliced generated sequences. The generated sequences are diverse, with 0% duplication rate at 1,000 sampled sequences and 4% duplication rate at 100,000 samples. Hexamer entropy = 8.31 out of 12 bits. (E) Validation of 1,000 generated sequences against the RNA-Seq measured MPRA using nearest neighbors. The first dense layer of the fitness predictor was used as feature space (256 features). Measured PSIs of the entire MPRA (black dots) are plotted with the interpolated PSIs of the generated sequences (yellow dots), estimated from 10 neighbors. Mean MPRA dPSI = 0.07 (+- 0.10). Mean dPSI of generated sequences = 0.38 (+- 0.06).

Next, we trained an exploration network with the same generator architecture as in **Figure 2C** to maximize the difference in predicted cell type-specific PSI between MCF7 and CHO (**Figure 5C**). We used the trained generator to sample 1,000 sequences, the majority of which were predicted by the neural net to be far more differentially spliced than any of the test sequences from the MPRA (**Figure 5D**; mean predicted dPSI of generated sequences = 0.56, compared to the average dPSI = 0.08 of the MPRA test set). For validation, we compared the generated sequences to the measured MPRA using a Nearest Neighbor search. We found that the DEN indeed learned to sample regulatory sequences centered on maximal differential splicing between the target cell lines (**Figure 5E, S5D**; mean NN-dPSI of generated sequences = 0.38, mean measured dPSI of MPRA sequences = 0.07).

Finally, we replicated the analysis using a linear logistic regression model with hexamer counts as features rather than a convolutional neural network fitness predictor. By reducing the regression model to a set of differentiable tensor operations, we could seamlessly integrate the model in the DEN pipeline (**Figure S5E-F**). Allowing both high and low-variance models enable users to better tailor the predictor properties for their given task. In some applications it may even be suitable to compose predictor ensambles to increase rigidity. In our case, we could re-train the DEN to jointly maximize the neural network and hexamer regression predictors, striking a balance between the two models (**Figure S5G**).

## DISCUSSION

We developed an end-to-end differentiable generative network architecture, Deep Exploration Networks (DENs), capable of synthesizing large, diverse sets of sequences with high fitness. The model could generate PASs which precisely conformed to target isoform ratios and 3’ cleavage positions, and even generated maximally used PASs that were far stronger than any previously designed sequence. Furthermore, the model could learn to generate maximally differentially spliced sequences.

DENs incorporate many techniques to improve its generative capabilities, but the single most important contribution was the control of exploration within the cost function during training. By having the generator sample two sequences given two random seeds, we developed a hinge-style loss which penalized sequence pairs that were similar above a certain threshold. Our analysis showed that the magnitude by which we punish sequence similarity (*repel weight*) almost entirely determines final generator diversity and, importantly, also largely determines the final fitness of the generated patterns. Taken as an expectation over all random input seeds sampled during training, the valley in the cost landscape to which our generator has access is enlarged by stochastically repelling similar sequences, and the expected width of the valley increases with a larger repel weight. During training, the optimizer trades off exploring (repelling similar patterns) with exploiting (maximizing pattern fitness) based on the temperature (repel weight) until convergence is reached. This scheme produces generative models which are (1) highly optimal, and (2) controllably diverse.

Another concern in sequence design is computational efficiency; for some applications, we may want to generate millions of candidate patterns, e.g. to synthesize in an oligo pool. Here, feed-forward models really outshine per-sequence optimization methods such as gradient ascent. In (Bogard et. al., 2019), it took roughly 2,000 updates (∼150 seconds on a CPU) to optimize a single sequence. In contrast, we train a DEN for 25,000 updates, i.e. more than a 10-fold increase. However, once training is done, the DEN encodes the distribution of sequences conditioned on the objective, enabling sampling from the distribution with a single feed-forward pass (∼0.010 seconds on a CPU). Hence, on a CPU with a trained DEN, we can generate 100,000 sequences in under 20 minutes, whereas it would take roughly 0.5 to 1 year to do it with gradient ascent.

In future work, there are several technical aspects to explore. First, the sequence similarity loss coefficient (repel weight) is currently kept constant. While this efficiently enforces exploration, it may in some cases be too rigid and require tuning before striking a good balance with the fitness objective. In particular, certain cost landscapes might have “pointy” – deep but narrow – valleys. With an overly large repel weight, neither of the generated sequences may ever be allowed down the valley before being pushed away by other repelling sequences. Rather, we would like to discover the pointy valleys during a high temperature stage of the training where exploration gradients are weighted more, and then descend at a later, low-temperature exploitation stage. We can easily adapt DENs to this scheme by treating the repel weight as a simulated annealing temperature which we decrementally lower throughout training, creating a “Deep Annealing”-style model.

We currently punish similarity by sequence-level comparison. But, depending on the application, the generated diversity may be sought in other metrics. We can easily generalize the similarity loss to allow any (differentiable) comparator function of the generated patterns, for example, we could penalize pattern similarity via a model of secondary structure. Another aspect is the method by which we pass the generated pattern to the predictor. Our results showed that dual input form representation, using both the continuous PWM and sampled one-hot patterns, can improve convergence. We approximated gradients with ST estimation, however, there may be more efficient estimators for multinomial sampling. In particular, training may improve using the Gumbel Softmax approximation (Jang et. al., 2016). Or, instead of using discrete samples, we may want to rectify the PWM representation. Specifically, it may be possible to re-train the deep learning predictor to give appropriate responses not only on 1-hot patterns, but also on arbitrary PWMs. During predictor model training, we can simultaneously feed the model randomly generated PWMs and enforce in the loss function that the model predicts the weighted average response of what it would predict for 1-hot patterns sampled from the PWM. We further showed that we could train “Inverse regression” models; a continuous-valued target was passed as input together with a random seed to sample sequences conforming to the target. By explicitly enforcing diversity, DENs are capable of generating multiple candidate samples (e.g. sequences) given one target input. Hence, they may be useful as deep probabilistic decoders. For example, we could construct a DEN which, given random input seeds and a single amino acid sequence, samples a diverse set of highly likely 3D protein structures.

We also showed in the case of differential splice sites that we could optimize generation for both a neural network and hexamer regression predictor. The hexamer regression model, by its low-variance design, provides regularization. However, we could provide more general regularization by inserting a customizable tensor model between the generator and predictor. For example, in concordance with (Killoran et. al, 2017), we may want to restrict generation to sequences that are conserved in the human genome. This could be achieved by placing a sequence GAN trained on the genome between the generator and predictor, such that the generator, given a random input seed, learns to generate optimal/high-fitness seeds for the GAN. The GAN in turn generates the final sequence passed to the predictor.

Experimental assays provide us with powerful tools to validate generative models, by enabling generated sequences to be synthesized and measured. Here, we tested a subset of our generated PASs. Remarkably, we observed that the new PASs were orders of magnitude stronger than any previously known sequence. While this performance owes to the power of deep exploration nets, it also owes to the consistent accuracy of the predictor models, even at the extremes of the regulatory distributions. In some applications, the predictor may not be sufficiently accurate, and the generated samples may reveal incorrect predictions once tested in the lab. Similar to earlier work on generative models employed for molecular design (Segler et. al., 2017), we envision DENs to be part of an adaptive sampling scheme, where the network generates a set of sequences which, after synthesis and measurements, provide augmented training data for the predictor, and this cycle is repeated until the generated patterns are in concordance with real biology.

Beyond splicing and 3’ end mRNA processing, there are many suitable biological applications for deep exploration networks. DENs could be used together with gene expression data to engineer cell type-specific enhancer or promoter sequences with differential affinities. DENs may also prove useful for generating candidate CRISPR Cas9 guide RNA which minimize predicted off-target effects (Lin et. al. 2018; Chuai et. al., 2018; Wang et. al., 2019). Moving on to the world of proteins, we imagine a vast number of application areas for deep exploration nets. Rational design of heterodimer pairs with orthogonal interaction has recently been demonstrated (Chen et. al., 2019). As more data is collected and used to train functional models of interaction, we would be able to use DENs for generation of candidate orthogonal binder sets, or even generalized interaction graphs. We could further combine DENs with existing neural network predictors to engineer protein function (Biswas et. al., 2018) or target structure (AlQuraishi et. al., 2019), or even use DENs as a means of sampling maximally likely structure predictions (Evans et. al., 2018). Finally, DENs could be utilized for generation of stable protein structures as well as high-specificity aptamer candidate sets.

## Acknowledgments

This work was supported by NIH R01HG009136 and R01HG009892 to G.S. We are grateful to Eric Klavins and Max Darnell for helpful comments on the manuscript.

---

**Figure S2.**
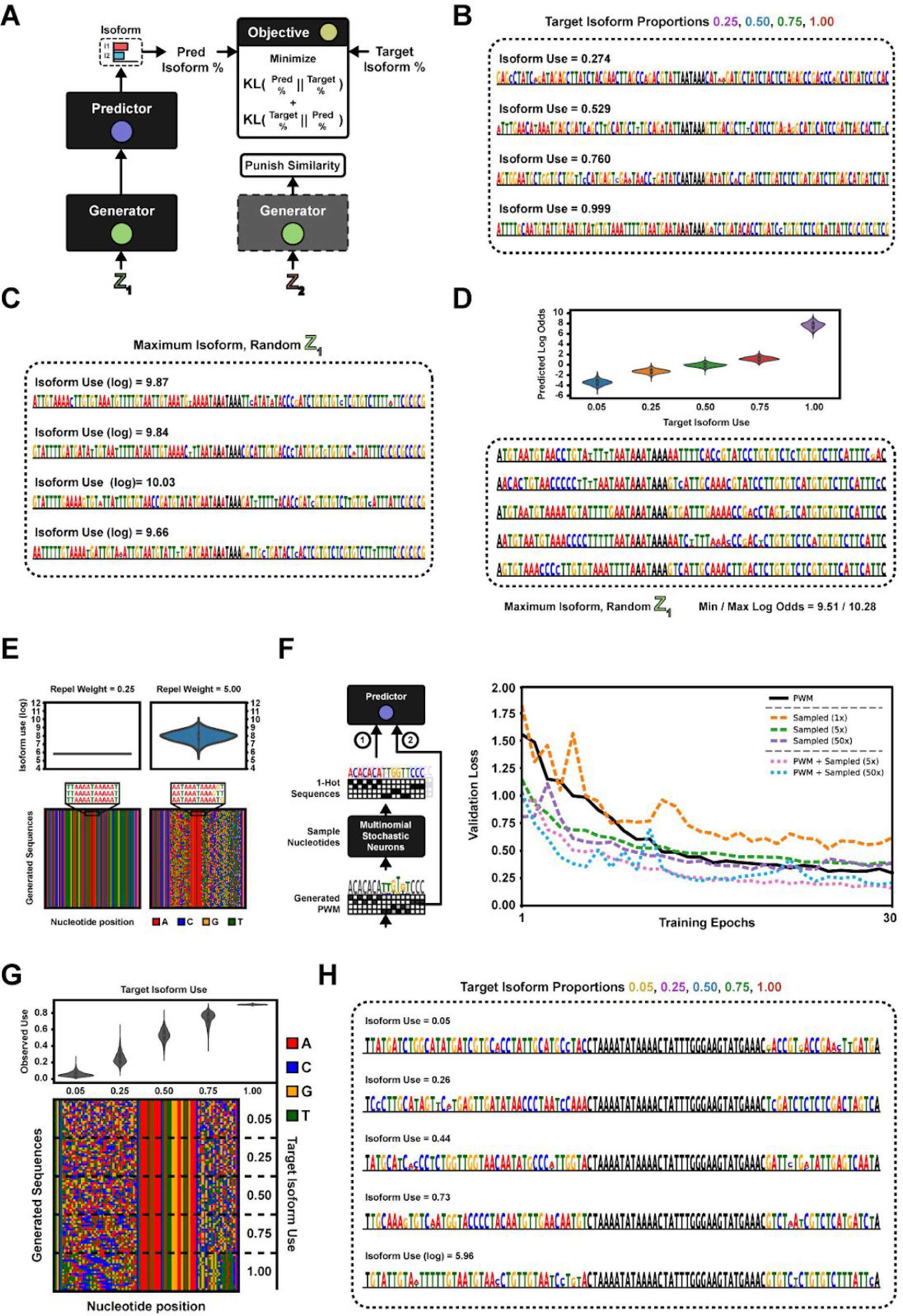
Generation Examples, Validation Experiments and Sequence Representations. Related to Figure 2. (A) The 1-hot sequence outputted by the generator is passed as input to the APA fitness predictor, which in turn outputs an isoform proportion prediction. This prediction is used in a symmetric KL-divergence loss to fit the generator to a fixed target isoform proportion. (B) Example sequences generated by four of the five deep exploration network instances evaluated in Figure 2D: The 25%, 50%, 75% and 100% (‘Max’) generators. (C) Example sequences generated by the 100%-target (‘Max’) APA isoform exploration network, using four different random seeds as input to the generator. (C) The entire analysis of (Figure 2A-E, S2A-C) was replicated using a different 3’ UTR sequence template (the fixed sequence regions). (Top) Predicted isoform proportions of 1,000 sampled sequences per target objective. Mean and Std dev of isoform log odds per target (proportions in parenthesis): (Target 5%) −3.40 +- 0.54 (= 3.70% +- 2.17%), (Target 25%) −1.26 +- 0.40 (= 22.9% +- 6.78%), (Target 50%) −0.064 +- 0.36 (= 48.5% +- 8.70%), (Target 75%) 1.13 +- 0.38 (75.0% +- 7.03%), (Target Max) 7.90 +- 0.68 (99.95% +- 0.04%). (Bottom) Five example sequences generated by the 100%-target (‘Max’) DEN, using random seeds as input to the generator. All five target APA generators are diverse, with 0% duplication rate at 100,000 sampled sequences by any of the generators. Hexamer entropy ranges between 8.87 and 9.20 bits depending on generator (of 12 bits maximum), with 1,970 to 2,591 unique hexamers across the first 1,000 sequences depending on generator. (E) Continuing the replication of (Figure 2A-E, S2A-B), the 100%-target (‘Max’) generator was re-trained twice, once with low similarity loss coefficient (repel weight; left) and in another instance a high coefficient (right). (Top) Predicted isoform proportions for 1,000 generated sequences, per generator instance. (Bottom) Sequence diversity illustrated by sampling 100 sequences. Low repel weight (left): Mean predicted isoform log odds = 5.78 +- 0.0, and 100% duplication rate at 100,000 sampled sequences. High repel weight (right): Mean isoform log odds = 7.64 +- 0.65, and 0% duplication rate at 100,000 samples. (F) Evaluation of three different sequence pattern representations: (1) Sampling a number of discrete 1-hot-coded patterns from the generated PWM and using as input the predictor, (2) Passing the generated PWM directly as input or (3) passing both representations as input to the predictor and walking down the average gradient. The methods were evaluated by training a deep exploration network to generate maximally used (100%-target) polyA signals, and tracking a validation loss across the first 30 epochs. Each epoch consists of 1,000 mini-batches of randomly generated input seeds. The validation loss is computed as the KL Divergence of the predicted isoform proportions compared against 100% usage on 500 random mini batches. Note, the validation loss is always computed using the consensus 1-hot-coded pattern extracted from the generated PWMs as input (guaranteeing well-formed input to the predictor). Reported are the median loss curves of five independent runs. (G) The deep exploration networks were validated against real RNA-Seq measurements using a nearest neighbor approximation. Five new generators were trained to produce polyA signals for the APA isoform targets 5%, 25%, 50%, 75%, 100%, but this time with a shorter (60 nt) freely tunable sequence so as to reduce “curse of dimensionality”-effects in the nearest neighbor search. TOMM5 APA reporter sequences were collected from (Bogard et. al., 2019) and the first dense layer activations predicted by APARENT were used as features when storing the sequences in a nearest neighbor database. Next, 1,000 sequences were generated from each of the five deep exploration nets, and their corresponding nearest neighbors were looked up. (Top) Measured isoform proportions (from RNA-Seq data) for the 50 nearest neighbors of each of the generated sequences. Mean and Std dev of isoform proportions: (Target 5%) 6.00% +- 3.03%, (Target 25%) 25.1% +- 6.95%, (Target 50%) 53.2% +- 7.05%, (Target 75%) 73.7% +- 7.18%, (Target Max) 88.05% +- 0.05%. (Bottom) Sequence diversity for each of the five generators, illustrated by sampling 20 sequences per target/generator and plotting the nucleotides on a 2-dimensional pixel grid. The first four target-isoform generators have 0% duplication rate at 100,000 sampled sequences, and the 100%-target (‘Max’) generator has a 0.1% duplication rate at 100,000 samples. Hexamer entropy ranges between 7.89 and 9.00 bits depending on generator (of 12 bits max), with 1,142 to 2,295 unique hexamers across the first 1,000 sequences depending on generator. (H) Example sequences generated by the five exploration networks trained in Figure S2G.

**Figure S3.**
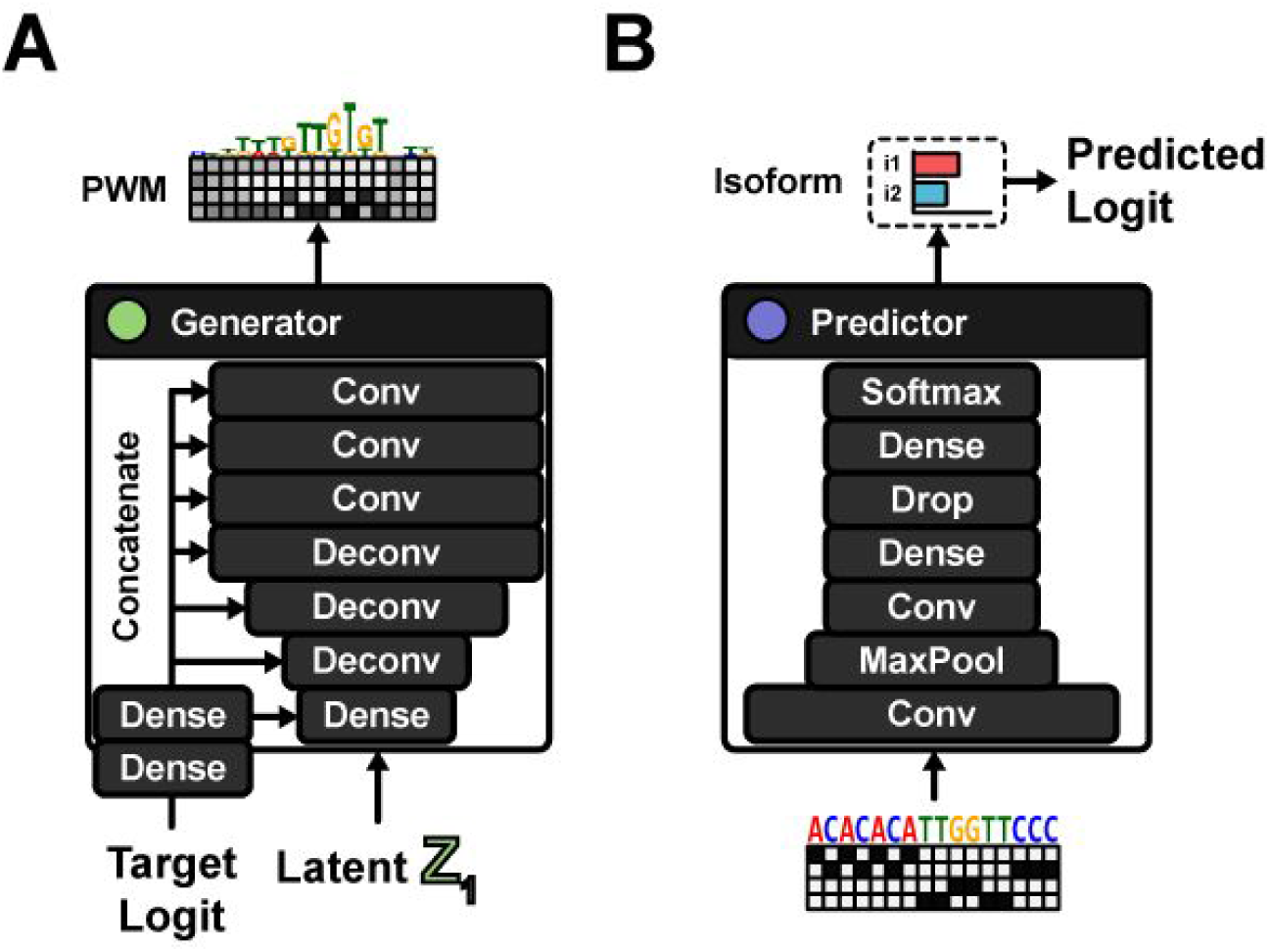
3’ Inverse APA Regression Generator and Predictor. Related to Figure 3. (A) A target isoform logit is passed as input to the generator in addition to the random seed. The logit is transformed into a high-dimensional, trainable embedding. The embedding is concatenated to the input tensor of every layer, enabling conditional learning and generation. (B) The APA fitness predictor architecture. A set of convolutional layers, pooling layers, dropout layers and dense layers transform the 1-hot-coded input sequence into an APA isoform proportion. The logit of the proportion is computed and passed to the objective.

**Figure S4.**
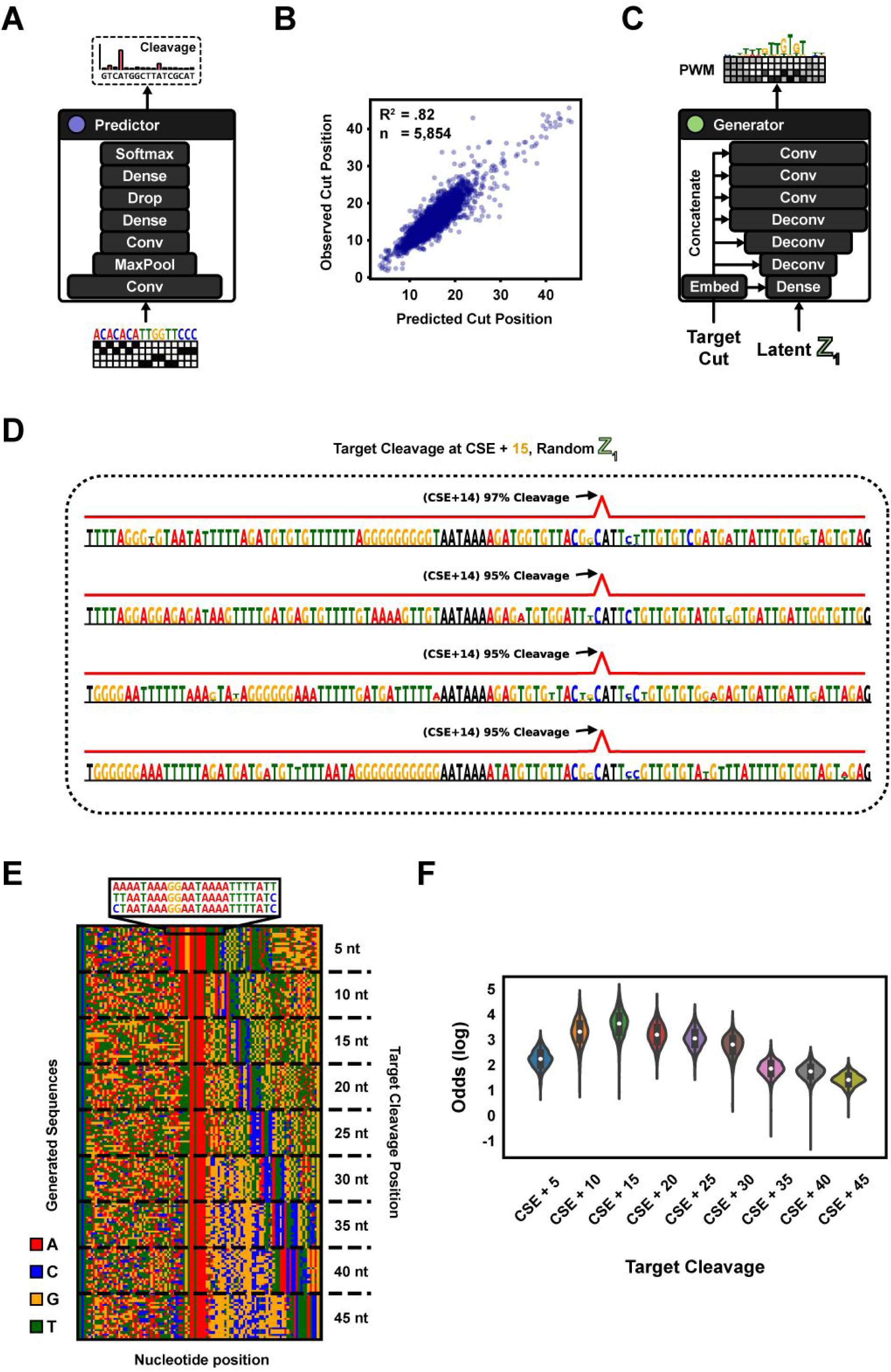
3’ Cleavage Predictor Model and Example Sequences. Related to Figure 4. (A) The 3’ cleavage predictor architecture. A set of convolutional layers, pooling layers, dropout layers and dense layers transform the 1-hot-coded input sequence into a 3’ cleavage distribution, predicting % Cleavage at nucleotide resolution. (B) Mean cut position prediction accuracy of the APARENT model. Shown are the predicted and measured mean cut positions of 5,854 test set sequences of the Alien2 APA reporter library from (Bogard et. al., 2019). Predicted vs. measured mean cut position R^2 = 0.82. (C) A target cut position (class index) is passed as input to the generator. The class index is transformed by a trainable embedding layer and concatenated onto every input tensor of every layer, enabling conditional generation. (D) Example sequences generated by the 3’ cleavage exploration network, using four different random seeds and the (+15 cut position)-class index as input to the generator. (E) Generator sequence diversity evaluated across all 9 target position classes, illustrated by randomly sampling 20 sequences per target position, and plotting them in a grid where rows denote sequences and columns denote nucleotide position. 0% duplication rate at 100,000 sampled sequences. Hexamer entropy = 9.07 of 12 bits, with 1,727 unique hexamers across the first 1,000 sequences. (F) Predicted cleavage log odds of 1,000 sampled sequences per target cleavage position. Mean and Standard deviation of cleavage log odds at each target cut position, in order: 2.10 (+- 0.34), 3.17 (+- 0.50), 3.54 (+- 0.47), 3.03 (+- 0.46), 2.93 (+- 0.38), 2.64 (+- 0.39), 1.73 (+- 0.32), 1.61 (+- 0.29), 1.31 (+- 0.24).

**Figure S5.**
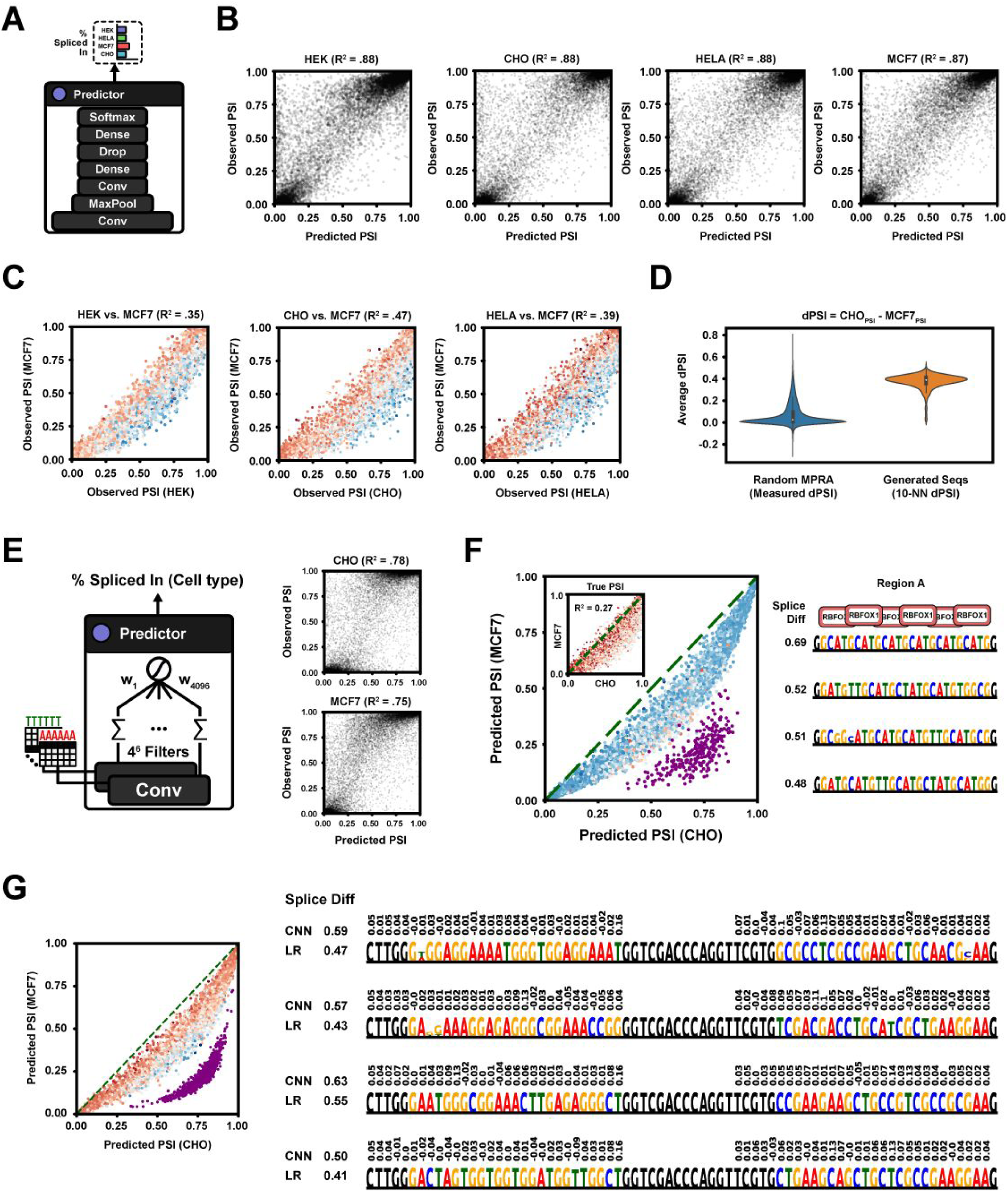
Cell Type-Specific Splicing Predictor Models and Sequence Generation Validation. Related to Figure 5. (A) The neural net predictor architecture. Convolutional, pooling, dropout and dense layers transform an input sequence into cell type-specific splice donor usage (PSI) predictions. (B) Predicted vs. measured PSI on a held-out test set of the splicing MPRA (n = ∼13,000). Predictions were made using a convolutional neural net with cell type-specific PSI outputs. (C) Comparison of measured PSIs between pairs of cell lines in the splicing MPRA test set (n = ∼13,000). Measured PSIs are displayed on the X/Y axes, and color intensity indicates the neural network-predicted dPSIs (blue/red = more/less PSI in cell type X than Y). (D) The max-differential splicing DEN was validated against real RNA-Seq measurements of the measured splicing MPRA using a nearest neighbor approximation. All ∼260,000 sequences of the splicing MPRA were transformed into 256-dimensional feature vectors, using the fitness predictor up until the first dense layer as a feature transform, and then stored in a nearest neighbor database. The feature transform reduces “curse of dimensionality”-effects in the nearest neighbor search and provides a degree of local motif invariance. Next, 1,000 sequences sampled from the DEN were transformed into 256-dimensional feature vectors according to the neural network predictor and looked up in the nearest neighbor database. The measured PSIs of the 10 nearest neighbors per generated sequence were used to estimate average PSIs. Shown are the measured and NN-interpolated MCF7-CHO dPSIs of the MPRA and generated sequences respectively. Mean MPRA dPSI = 0.07 (+- 0.10). Mean dPSI of generated sequences = 0.38 (+- 0.06). (E) A differentiable relaxation of a 6-mer logistic regression model. A convolutional layer with 4096 filters, each encoding a distinct 6-mer (filter weight = 6, bias = −5), result in 4096 activation maps which after a summation over positions become hexamer counts. The counts are combined with cell type-specific weights and squashed through a sigmoid. (F) Evaluation of a deep exploration network, using a hexamer regression predictor, when tasked with generating 500 maximally differentially spliced sequences between cell lines MCF7 and CHO. (Left) Percent Spliced In (Splice donor usage) as predicted by the regression model on all MPRA test set sequences in MCF7 and CHO. Color intensity indicates measured dPSI. Purple dots indicate the 500 generated sequences. (Inline) Inverted scatter plot; Measured PSI between cell types for all MPRA test sequences are shown on the axes and color indicates predicted dPSI. Measured vs. Predicted dPSI R^2 on test set = 0.27. (Right) Selection of maximally differentially spliced generated sequences. The generator has learned to exploit the additive independence of hexamer scores, by populating the sequences with a very differential motif, GCATGC (RBFOX1 binding site). (G) Both the neural network and hexamer regression splicing models were used as fitness predictors, and the exploration network was retrained to maximize differential splicing between MCF7 and CHO according to the average response of both predictors. The generator was used to sample 1,000 sequences, and their average predicted PSIs (purple) were plotted alongside the predicted PSIs of the splicing MPRA test set (color intensity indicates measured dPSI). Mean predicted dPSI of test sequences = 0.08 (+- 0.07). Mean predicted dPSI of generated sequences = 0.54 (+- 0.04). (Right) The top four differentially spliced sequences generated by the DEN are shown with the neural net (CNN) and hexamer regression (LR) dPSI predictions. Shown above each sequence are the hexamer weights.

**Movie S1. Random Sampling of Sequences during 3’ Cleavage DEN Training. Related to Figure 4.**

Evaluation of 20 randomly generated sequences per target cut objective by continuously sampling new input seeds during DEN training. For each generator weight update (each mini batch of seeds), 20 uniformly random seeds per cut objective are passed to the generator. The corresponding generated sequences are updated on a 2-dimensional pixel grid where nucleotides are represented as different colors (A = red, C = blue, G = orange, T = green). The loss curve of each generated sequence is simultaneously tracked in real-time. One of the 20 randomly generated sequences is displayed to the right for each cut objective. The black-colored nucleotides in the PWMs are fixed (non-changeable) sequence regions. We fixed the CSE (‘AATAAA’) and target cut dinucleotide during optimization.

**Movie S2. Tracking a Fixed Sequence Set during 3’ Cleavage DEN Training. Related to Figure 4.**

Continuous evaluation of 20 fixed generator input seeds throughout 3’ Cleavage DEN training. For each weight update, the generator is tasked with generating the sequence set for a fixed set of seeds. The generated sequences are tracked through time during training and updated on a 2-dimensional pixel grid where nucleotides are represented as different colors (A = red, C = blue, G = orange, T = green). The loss curve of each generated sequence is simultaneously tracked in real-time. One of the generated sequences is displayed to the right for each cut objective. The black-colored nucleotides in the PWMs are fixed (non-changeable) sequence regions. We fixed the CSE (‘AATAAA’) and target cut dinucleotide during optimization.

## REFERENCES

Abadi, M., Agarwal, A., Barham, P., Brevdo, E., Chen, Z., Citro, C., … & Ghemawat, S. (2016). Tensorflow: Large-scale machine learning on heterogeneous distributed systems (arXiv).

Alipanahi, B., Delong, A., Weirauch, M.T., and Frey, B.J. (2015). Predicting the sequence specificities of DNA- and RNA-binding proteins by deep learning. Nature Biotechnology 33, 831–838.

AlQuraishi, M. (2019). End-to-end differentiable learning of protein structure. Cell Systems 8, 292–301.

Avsec, Ž., Weilert, M., Shrikumar, A., Alexandari, A., Krueger, S., Dalal, K., … & Zeitlinger, J. (2019). Deep learning at base-resolution reveals motif syntax of the cis-regulatory code (bioRxiv).

Avsec, Ž., Kreuzhuber, R., Israeli, J., Xu, N., Cheng, J., Shrikumar, A., … Gagneur, J. (2019). The Kipoi repository accelerates community exchange and reuse of predictive models for genomics. Nature Biotechnology 37, 592–600.

Bengio, Y., Léonard, N., & Courville, A. (2013). Estimating or propagating gradients through stochastic neurons for conditional computation (arXiv).

Biswas, S., Kuznetsov, G., Ogden, P. J., Conway, N. J., Adams, R. P., & Church, G. M. (2018). Toward machine-guided design of proteins (bioRxiv).

Blencowe, B. J. (2006). Alternative splicing: new insights from global analyses. Cell 126, 37–47.

Bogard, N., Linder, J., Rosenberg, A. B., & Seelig, G. (2019). A Deep Neural Network for Predicting and Engineering Alternative Polyadenylation. Cell 178, 91–106.

Brookes, D. H., Park, H., & Listgarten, J. (2019). Conditioning by adaptive sampling for robust design (arXiv).

Chen, Z., Boyken, S. E., Jia, M., Busch, F., Flores-Solis, D., Bick, M. J., … Baker, D. (2019). Programmable design of orthogonal protein heterodimers. Nature 565, 106–111.

Chollet, F. (2015). Keras.

Chuai, G., Ma, H., Yan, J., Chen, M., Hong, N., Xue, D., … Liu, Q. (2018). DeepCRISPR: optimized CRISPR guide RNA design by deep learning. Genome Biology 19, 80.

Courbariaux, M., Hubara, I., Soudry, D., El-Yaniv, R., & Bengio, Y. (2016). Binarized neural networks: Training deep neural networks with weights and activations constrained to+ 1 or-1 (arXiv).

Cuperus, J. T., Groves, B., Kuchina, A., Rosenberg, A. B., Jojic, N., Fields, S., & Seelig, G. (2017). Deep learning of the regulatory grammar of yeast 5′ untranslated regions from 500,000 random sequences. Genome Research 27, 2015–2024.

Di Giammartino, D. C., Nishida, K., & Manley, J. L. (2011). Mechanisms and consequences of alternative polyadenylation. Molecular Cell 43, 853–866.

Eiben, A. E., & Smith, J. (2015). From evolutionary computation to the evolution of things. Nature 521, 476–482.

Elkon, R., Ugalde, A. P., & Agami, R. (2013). Alternative cleavage and polyadenylation: extent, regulation and function. Nature Reviews Genetics 14, 496–506.

Evans, R., Jumper, J., Kirkpatrick, J., Sifre, L., Green, T. F. G., Qin, C., … & Petersen, S. (2018). De novo structure prediction with deeplearning based scoring. Annual Reviews of Biochemistry 77, 363–382.

Eraslan, G., Avsec, Ž., Gagneur, J., & Theis, F. J. (2019). Deep learning: new computational modelling techniques for genomics. Nature Reviews Genetics 20, 389–403.

Goodfellow, I., Pouget-Abadie, J., Mirza, M., Xu, B., Warde-Farley, D., Ozair, S., … & Bengio, Y. (2014). Generative adversarial nets. In 2014 Advances in neural information processing systems, 2672–2680.

Greenside, P., Shimko, T., Fordyce, P., & Kundaje, A. (2018). Discovering epistatic feature interactions from neural network models of regulatory DNA sequences. Bioinformatics 34, 629–637.

Jaganathan, K., Kyriazopoulou Panagiotopoulou, S., McRae, J. F., Darbandi, S. F., Knowles, D., Li, Y. I., … Farh, K. K.-H. (2019). Predicting Splicing from Primary Sequence with Deep Learning. Cell 176, 535–548.

Jang, E., Gu, S., & Poole, B. (2016). Categorical reparameterization with gumbel-softmax (arXiv).

Kelley, D. R., Snoek, J., & Rinn, J. L. (2016). Basset: learning the regulatory code of the accessible genome with deep convolutional neural networks. Genome Research 26, 990–999.

Kelley, D. R., Reshef, Y. A., Bileschi, M., Belanger, D., McLean, C. Y., & Snoek, J. (2018). Sequential regulatory activity prediction across chromosomes with convolutional neural networks. Genome Research 28, 739–750.

Killoran, N., Lee, L. J., Delong, A., Duvenaud, D., & Frey, B. J. (2017). Generating and designing DNA with deep generative models (arXiv).

Kingma, D. P., & Ba, J. (2014). Adam: A method for stochastic optimization (arXiv).

Kingma, D. P., & Welling, M. (2013). Auto-encoding variational bayes (arXiv).

Lanchantin, J., Singh, R., Lin, Z., & Qi, Y. (2016). Deep motif: Visualizing genomic sequence classifications (arXiv).

Lee, Y., & Rio, D. C. (2015). Mechanisms and regulation of alternative pre-mRNA splicing. Annual Reviews of Biochemistry 84, 291–323.

Lin, J., & Wong, K. C. (2018). Off-target predictions in CRISPR-Cas9 gene editing using deep learning. Bioinformatics 34, 656–663.

Maaten, L. V. D., & Hinton, G. (2008). Visualizing data using t-SNE. Journal of Machine Learning Research 9, 2579–2605.

Mirjalili, S., Dong, J. S., Sadiq, A. S., & Faris, H. (2020). Genetic Algorithm: Theory, Literature Review, and Application in Image Reconstruction. Nature-Inspired Optimizers, Springer, Cham., 69–85.

Mirza, M., & Osindero, S. (2014). Conditional generative adversarial nets (arXiv).

Pitis, Silviu. (2017). Beyond Binary: Ternary and One-hot Neurons. Blog post on the R2RT blog. (Online) https://r2rt.com/beyond-binary-ternary-and-one-hot-neurons.

Quang, D., & Xie, X. (2019). FactorNet: A deep learning framework for predicting cell type specific transcription factor binding from nucleotide-resolution sequential data. Methods 166, 40–47.

Radford, A., Metz, L., & Chintala, S. (2015). Unsupervised representation learning with deep convolutional generative adversarial networks (arXiv).

Roca, X., Krainer, A. R., & Eperon, I. C. (2013). Pick one, but be quick: 5’ splice sites and the problems of too many choices. Genes & Development 27, 129–144.

Rocklin, G. J., Chidyausiku, T. M., Goreshnik, I., Ford, A., Houliston, S., Lemak, A., … & Arrowsmith, C. H. (2017). Global analysis of protein folding using massively parallel design, synthesis, and testing. Science 357, 168–175.

Rosenberg, A. B., Patwardhan, R. P., Shendure, J., & Seelig, G. (2015). Learning the sequence determinants of alternative splicing from millions of random sequences. Cell 163, 698–711.

Sample, P. J., Wang, B., Reid, D. W., Presnyak, V., McFadyen, I. J., Morris, D. R., & Seelig, G. (2019). Human 5’ UTR design and variant effect prediction from a massively parallel translation assay. Nature Biotechnology 37, 803–809.

Segler, M. H., Kogej, T., Tyrchan, C., & Waller, M. P. (2017). Generating focused molecule libraries for drug discovery with recurrent neural networks. ACS central science, 4, 120–131.

Shukla, A., Pandey, H. M., & Mehrotra, D. (2015). Comparative review of selection techniques in genetic algorithm. In 2015 International Conference on Futuristic Trends on Computational Analysis and Knowledge Management, IEEE, 515–519.

Simonyan, K., Vedaldi, A., & Zisserman, A. (2013). Deep inside convolutional networks: Visualising image classification models and saliency maps (arXiv).

Stewart, K., Chen, Y. J., Ward, D., Liu, X., Seelig, G., Strauss, K., & Ceze, L. (2018). A content-addressable DNA database with learned sequence encodings. In International Conference on DNA Computing and Molecular Programming, Springer, Cham., 55–70.

Tian, B., & Manley, J. L. (2017). Alternative polyadenylation of mRNA precursors. Nature Reviews. Molecular Cell Biology 18, 18–30.

Wang, D., Zhang, C., Wang, B., Li, B., Wang, Q., Liu, D., … & Wang, Y. (2019). Optimized CRISPR guide RNA design for two high-fidelity Cas9 variants by deep learning. Nature Communications 10, 1–14.

Zhou, J., & Troyanskaya, O. G. (2015). Predicting effects of noncoding variants with deep learning–based sequence model. Nature Methods 12, 931–934.

