## Supplementary Information for "Deep exploration networks for rapid engineering of functional DNA sequences"

### CONTACT FOR REAGENT AND RESOURCE SHARING

### EXPERIMENTAL MODEL AND SUBJECT DETAILS

Four cell lines were used in this study: Human embryonic kidney cells (HEK293; female), human cervical cancer cells (HeLa; female), human breast cancer cells (MCF7; female), and chinese hamster ovary cells (CHO; female).

### METHOD DETAILS

Here we present both the computational and experimental methods developed and applied in this paper. We start by describing the general mathematical framework of Deep Exploration Networks: The generator architecture, how sequence patterns are represented, pattern masking operations, integration of predictor models, and finally how the cost function is defined.

Next, we describe the three genomics applications considered in this paper, which fall into two categories: The polyadenylation part, and the splicing part. For each of these parts, we describe the predictor model used, the data it was trained on and the specific generator architecture and hyperparameter configuration used when training the DENs. We also describe the details of the physical experiments carried out to validate the performance of our generated sequences. In the case of APA, we describe the experiment for comparing DEN-generated Max-isoform signals to the baseline Gradient ascent-generated sequences, and we prove that the Ct-values measured in the qPCR assay can be used to calculate isoform odds ratio lower bounds.

First, we present the mathematical notation and style used throughout this text, as well as provide a list of important definitions.

**Math Notation**

Scalars are denoted as lowercase italic letters, e.g. $c$.

Vectors are denoted as lowercase italic letters with an arrow, e.g. $\vec{v}$.

Vector elements are referred to using square brackets and one subscript: $[v]_i$.

Matrices or higher-order tensors are denoted as uppercase letters, e.g. $A$.

Tensor elements are referred to using square brackets and multiple subscripts: $[a]_{ij}$.

**Definitions**

The list below briefly summarizes the most important variables and entities defined in detail in the section "Deep Exploration Network (DEN)" below. Note that there sometimes exist multiple independent instances of some of these variables, and we distinguish between variable instances using superscripts encased in parentheses (for example: $X^{(1)}$ and $X^{(2)}$).

$\mathcal{G}$ - Generator model function, i.e. $X = \mathcal{G}(\vec{z})$
$\vec{z}$ - Generator seed input
$X$ - Generator output matrix (nucleotide log probabilities)
$\mathcal{P}$ - Predictor model function, i.e. $\bar{y} = \mathcal{P}(P)$ or $y = \mathcal{P}(S)$
$P$ - PWM matrix obtained by normalizing $X$
$S$ - 1-hot-coded pattern sampled from $P$
$\bar{y}$ - Predictor output, obtained by running $\mathcal{P}$ on $P$
$y$ - Predictor output, obtained by running $\mathcal{P}$ on $S$

**Modeling & Optimization Software**

We used the auto-differentiation and deep learning package Keras in Python for all neural network implementations and training (Chollet, 2015). For training the generator networks, we used the Adam optimizer (Kingma et. al., 2014). To implement certain operations required during DEN training, we had to write custom code in Tensorflow (Abadi et. al, 2016). The Tensorflow implementation for multinomial straight-through gradient estimation (the 1-hot sampling operation for sequence PWMs) was based on (Silviu, 2017).

#### Deep Exploration Network (DEN)

Here we present the model, cost function and training procedure of Deep Exploration Networks. We describe the common, architecture here. For details regarding generator or predictor details, we refer the reader to the sections of the corresponding applications further below.

**Generator Architecture**

The generator model $\mathcal{G}$ is a feed-forward neural network which receives a seed vector $\vec{z} \in \mathcal{R}^D$ as input and outputs a pattern $X \in \mathcal{R}^{N \times M}$ according to $X = \mathcal{G}(\vec{z})$. Here N denotes the number of letters of the generated sequence pattern and M denotes the number of channels (the alphabet size). In the context of genomics, the alphabet is the set of nucleotides and M = 4. The generated pattern $X$ is treated as a matrix of nucleotide log probabilities. For some applications presented in this paper, the generator model takes auxiliary information as input in order to produce class-conditional patterns, in which case the formula can be described as $X = \mathcal{G}(\vec{z}, c)$, where $c \in \mathcal{N}$ is an integer representing the class index.

The exact architecture of the generator network changes depending on the application, but the overall design remains the same throughout the paper and is based on Deep Convolutional GANs (DC-GANs) (Radford et. al., 2015; See Figure 2C for a high-level illustration of the model). First, a dense layer with ReLU activations transform the input seed $\vec{z} \in \mathcal{R}^D$ into a high-dimensional vector. This vector is reshaped into a two-dimensional matrix, where the first dimension encodes sequence position and the second dimension encodes sequence channel. The position dimension is initially scaled down to a fraction of the final target sequence length, such that it can be upsampled by strided deconvolutions by subsequent layers. The channel dimension starts with a large number of channels (384 for all our models) and is gradually compressed to smaller numbers throughout the generator network, until the final layer outputs a sequence with only 4 channels (4 nucleotides). The dense layer is followed by a number of de-convolutional layers (3 layers in all our models). These layers are always configured to do strided deconvolution with an upsampling factor of 2, with ReLU activations. The filters vary in width between 6 and 8; we always perform valid convolutions in all our models, so the widths must be set to specific values such that the final generated sequence is of intended length. The de-convolutional layers are followed by convolutional layers (3 layers in all our models). All convolutional layers have a filter width of 8, a stride of 1, no pooling, and perform padded rather than valid convolutions (i.e. they do not change the sequence length). All convolutional layers except the final layer have ReLU activations; the final layer has linear activations, corresponding to nucleotide log probabilities. The final layer has 4 output/filter channels, corresponding to the 4 nucleotides. Between every de-convolutional or convolutional layer and its activation function (except the final layer), we perform batch normalization over the channel axis.

When training Deep Exploration Networks (DENs), two random seed vectors $\vec{z}^{(1)} \in \mathcal{R}^D$ and are independently sampled from uniform distributions during each forward pass:

$$\vec{z}_i^{(1)}, \vec{z}_i^{(2)} \in \mathrm{Unif}(-1, 1), \text{ for } i = 1, ..., D$$

The seeds are separately passed to and computed by the generator network, producing two output patterns (nucleotide log probability matrices) $X^{(1)} \in \mathcal{R}^{N \times M}$ and $X^{(2)} \in \mathcal{R}^{N \times M}$:

$$X^{(1)} = \mathcal{G}(\vec{z}_i^{(1)}), X^{(2)} = \mathcal{G}(\vec{z}_i^{(2)})$$

Gradients $\nabla_{W_G} X^{(1)} = \nabla_{W_G} \mathcal{G}(\vec{z}_i^{(1)})$ and $\nabla_{W_G} X^{(2)} = \nabla_{W_G} \mathcal{G}(\vec{z}_i^{(2)})$ with respect to the generator weights $W_G$ can be computed using standard differentiation in Keras, as the generator is a regular (de-)convolutional ReLU network.

The exact generator network configuration and architecture, including hyper parameter settings, differ for each of the three applications considered in this paper, and we refer the reader to their corresponding sections below for a detailed specification of each generator.

**Pattern Representation**

During training, we use two different representations of the generated patterns as input to the downstream fitness predictor model. The first pattern representation is referred to in this paper as RIFR (Relaxed Input Form Representation), and is equivalent to a sequence PWM (Position Weight Matrix). We apply a nucleotide-wise Softmax function to the generated matrices $X^{(1)}$ and $X^{(2)}$ to transform them into the corresponding PWMs $P^{(1)}$ and $P^{(2)}$ (Killoran et. al., 2017):

$$[p^{(1)}]_{ij} = \frac{e^{[x^{(1)}]_{ij}}}{\sum_{k=1}^{M} e^{[x^{(1)}]_{ik}}}, [p^{(2)}]_{ij} = \frac{e^{[x^{(2)}]_{ij}}}{\sum_{k=1}^{M} e^{[x^{(2)}]_{ik}}}$$

The Softmax function is differentiable, which means it is straight-forward to use an auto-differentiation package like Keras to compute the gradients $\nabla_{W_G} P^{(1)}$ and $\nabla_{W_G} P^{(2)}$.

The second pattern representation is referred to as SIFR (Sampled Input Form Representation) and consists of K discrete, 1-hot-coded samples drawn from the PWMs, where each nucleotide position corresponds to an independent multinomial distribution and the nucleotide weights at that position in the PWM correspond to the sampling probabilities:

$$J_{ik}^{(1)} \in \text{Multinom}([p^{(1)}]_{i1}, ..., [p^{(1)}]_{iM}), \text{ for } k = 1, ..., K$$

Here $J_{ik}^{(1)}$ is a random categorical variable that, in the case of DNA, can take on 4 different indices corresponding to the 4 different nucleotides. After sampling a nucleotide index at each position in the sequence, we construct K one-hot-coded matrix samples $S_k^{(1)}$:

$$[s^{(1)}]_{kij} = \begin{cases} 1, & \text{if } j = J_{ik}^{(1)} \\ 0, & else \end{cases}$$

We similarity sample nucleotide indices and construct matrices $S_k^{(2)}$ from the second PWM.

$$J_{ik}^{(2)} \in \text{Multinom}([p^{(2)}]_{i1}, ..., [p^{(2)}]_{iM}), \text{ for } k = 1, ..., K$$

$$[s^{(2)}]_{kij} = \begin{cases} 1, & \text{if } j = J_{ik}^{(2)} \\ 0, & else \end{cases}$$

We employ straight-through (ST) estimation to approximate gradients $\nabla_{W_G} S^{(1)}$ and $\nabla_{W_G} S^{(2)}$ and implement custom operations in Keras/Tensorflow to compute the ST gradients during generator training (Bengio, Léonard & Courville, 2013; Courbariaux et. al., 2016; Bogard et. al., 2019; Silviu, 2017).

**Sequence Masks & Templates**

In certain scenarios, the user may not always want the entire sequence pattern to be randomly initialized and changeable by the generator. Depending on the predictor model's input window, the user may need to pad the changeable sequence with fixed sequence, and in some cases the user may want to bias the generation by locking particular motifs in the sequence.

To provide support for this, we define a mask matrix, $M \in \{0,1\}^{N \times M}$, where we encode 1's at changeable nucleotide positions and 0's at fixed positions:

$$[m]_{ij} = \begin{cases} 1, & \text{if } i \text{ is not fixed} \\ 0, & else \end{cases}$$

We also define a template matrix, $T \in \{0,1\}^{N \times M}$, where we encode 0's at changeable nucleotide positions and a 1 at every fixed position and column corresponding to the specific nucleotide identity at said position:

$$[t]_{ij} = \begin{cases} 1, & \text{if } i \text{ is fixed and } j \text{ encodes the nucleotide at position } i \\ 0, & else \end{cases}$$

Let us rename the PWMs $P^{(1)}, P^{(2)}$ defined above (before masking and templating) as $P^{(1)}_{init}, P^{(2)}_{init}$ respectively. We then re-define $P^{(1)}, P^{(2)}$ as:

$P^{(1)} = P^{(1)}_{init} \times M + T$ and $P^{(2)} = P^{(2)}_{init} \times M + T$

Here we multiplied the PWM by the mask matrix $M$, effectively zeroing out all elements corresponding to fixed positions (giving them zero gradients), and adding the template matrix $T$ , which adds a one-hot-coded sub-pattern over fixed positions while still propagating zero gradients (due to the prior masking operation). Note that the gradients $\nabla_{W_G} P^{(1)}$ and $\nabla_{W_G} P^{(2)}$ are still well-defined and calculable by auto-differentiation; they are zero for masked (fixed) nucleotide positions and equal to $\nabla_{W_G} P^{(1)}_{init}$ and $\nabla_{W_G} P^{(2)}_{init}$ for non-masked positions.

Sample patterns $S^{(1)}_k$ and $S^{(2)}_k$ are still defined in terms of $P^{(1)}$ and $P^{(2)}$ as before.

**Predictor Integration**

Any differentiable predictor model $\mathcal{P}$ is compatible with Deep Exploration Networks, i.e. if the predictor forward pass is formulated as $y = P(X)$, where $X$ is the input tensor and $y$ is the model prediction, then the gradient $\nabla_X P(X)$ must be defined (Simonyan et. al., 2013; Lanchantin et. al., 2016).

In the case of RIFR, we directly pass one of the two generated PWMs ($P^{(1)}$) to compute the prediction output, $\bar{y}^{(1)} = \mathcal{P}(P^{(1)})$. In the case of SIFR, we separately pass all of the K 1-hot-coded samples ($S_k^{(1)}, \forall k$) to the predictor model, $y_k^{(1)} = \mathcal{P}(S_k^{(1)}), \text{ for } k = 1, ..., K$.

Using the chain rule, and the assumption that $\nabla_X P(X)$ is always defined, we can automatically compute the gradients $\nabla_{W_G} \bar{y}^{(1)}$ and $\nabla_{W_G} y_k^{(1)}, \forall k$ using Keras.

**Cost Function & Training**

We typically minimize four distinct cost terms when training a DEN: (1) The fitness objective, (2) the similarity loss, (3) the entropy and, optionally, (4) any sequence regularization. The general formula of the cost function can be written as:

$$C = C_{\text{objective}} + C_{\text{regularization}} + C_{\text{entropy}} + C_{\text{similarity}}$$

***Cost term:*** $C_{\text{objective}}$

The fitness cost $C_{\text{objective}}$ defines the biological target of the sequence pattern. In RIFR, it is a function of the continuous PWM predictor output $\bar{y}^{(1)}$, i.e. $C_{\text{objective}} = C_{\text{objective}}(\bar{y}^{(1)})$. In SIFR, it is a function of the K sampled prediction outputs $y_k^{(1)}, \forall k$: $C_{\text{objective}} = C_{\text{objective}}(y_k^{(1)})$. And finally, in DIFR, $C_{\text{objective}}$ is the average of the two losses, using both the PWM and sample-predictions: $C_{\text{objective}} = C_{\text{objective}}(\bar{y}^{(1)}, y_k^{(1)}) = (C_{\text{objective}}(\bar{y}^{(1)}) + C_{\text{objective}}(y_k^{(1)}))/2$. Exactly which loss function is used depends entirely on the application. In the case of differential expression, $C_{\text{objective}}$ may be a sum-of-squares error between predicted and target delta enrichment. In the case of isoform engineering, the loss is usually defined with KL-divergence. Any loss is valid as long as it maintains differentiability w.r.t. $\bar{y}^{(1)}$ and $y_k^{(1)}$.

***Cost term:*** $C_{\text{similarity}}$

The similarity cost should be a differentiable comparator function of the two generated patterns $S_k^{(1)}$ and $S_k^{(2)}$ (the sampled 1-hot-coded patterns) or $P^{(1)}$ and $P^{(2)}$ (the continuous PWMs), i.e. $C_{\text{similarity}} = C_{\text{similarity}}(S_k^{(1)}, S_k^{(2)})$ or $C_{\text{similarity}} = C_{\text{similarity}}(P^{(1)}, P^{(2)})$. Using the $C_{\text{similarity}}(S_k^{(1)}, S_k^{(2)})$-representation is similar in concept to applying a sparse L1-penalty directly on the PWMs; if two sampled elements are equal under the comparator, we penalize the patterns with a constant value (1 in the case of cosine similarity). Using $C_{\text{similarity}} = C_{\text{similarity}}(P^{(1)}, P^{(2)})$ corresponds to an L2-style penalty, where a comparator such as cosine similarity would punish two elements proportional to the magnitude of their PWM entries. For the specific applications considered in this paper, we use the sampled (L1) representation.

We use cosine similarity as the basis for our similarity penalty throughout the paper, defined in terms of $S_k^{(1)}$ and $S_k^{(2)}$. We found empirically that we get better results using a slack-bound margin, which allows a fraction of the nucleotides to be identical without incurring a penalty. We typically use a margin of 0.4, allowing up to 40% of sequences to share nucleotide content. Also, because we draw K independent 1-hot-coded samples per forward pass, the penalty is actually a mean cosine similarity metric. The cost function is thus defined as:

$$C_{\text{similarity}} = \frac{1}{K}\sum_{k=1}^{K}\max\left(\left[\frac{1}{N}\sum_{i=1}^{N}\sum_{j=1}^{M}[s^{(1)}]_{kij}\cdot[s^{(2)}]_{kij}\right]-\text{margin},0\right)$$

We noted that the DEN sometimes learned to generate sequence patterns which were offset by one or two nucleotides, effectively escaping the cosine similarity penalty. To rectify this, we developed a more rigorous penalty by comparing the two sequence patterns at different offsets:

$$C_{\text{offset}} = \max\left(\left[\frac{1}{N}\max_{\sigma}\sum_{i=1}^{N-\sigma}\sum_{j=1}^{M}[s^{(1)}]_{k,i+\max(\sigma,0),j}\cdot[s^{(2)}]_{k,i+\max(-\sigma,0),j}\right]-\epsilon,0\right)$$

Here, $\sigma = -\sigma_{max}$ to $\sigma_{max}$ defines the pattern offset (in discrete nucleotides), and we penalize the patterns by the maximum cosine similarity across all possible offsets. Theoretically, we should set $\sigma_{max} = N-1$ to test all possible offsets, however practically we found that $\sigma_{max} = 1$ was enough to remove generator offset artifacts.

***Cost term:*** $C_{\text{entropy}}$

We explicitly control the entropy of the generated PWMs by optimizing the negative Shannon Entropy for a target value. $C_{\text{entropy}}$ is thus a function of $P^{(1)}$, i.e. $C_{\text{entropy}} = C_{\text{entropy}}(P^{(1)})$, and since Shannon Entropy is differentiable we can compute the gradients $\nabla_{W_G}C_{\text{entropy}}(P^{(1)})$ w.r.t. the generator weights $W_G$ to train them. Empirically, we found that minimizing an absolute error between the average nucleotide entropy and a target entropy (in bits) works well:

$$C_{\text{entropy}} = \left(t_{\text{bits}} - \frac{1}{N}\sum_{i=1}^{N}\left(2-\sum_{j=1}^{M}-[p^{(1)}]_{ij}\cdot\log_2[p^{(1)}]_{ij}\right)\right)^2$$

***Cost term:*** $C_{\text{regularization}}$

We optionally penalize or reward specific motifs or nucleotide content in the generated pattern by shifted multiplication with itself to mask out sequence sub-patterns. This is useful for example to repress known artifacts of the predictor. Same as the similarity cost, we can define this cost either in terms of $S_k^{(1)}$ (the sampled 1-hot-coded patterns) or $P^{(1)}$ (the continuous PWM), i.e.

$C_{\text{regularization}} = C_{\text{regularization}}(S_k^{(1)})$ or $C_{\text{regularization}} = C_{\text{regularization}}(P^{(1)})$. We find that the sample-form ($S_k^{(1)}$), being a sparse L1-style loss, gives better results when penalizing motifs and the continuous form ($P^{(1)}$), being more similar to an L2 loss, gives better convergence for promoting particular motifs.

For each 1-hot-coded motif $F$ of length L, we either add or subtract the following component to/from $C_{\text{regularization}}$, to penalize or promote generation of $F$ respectively on the basis of $P^{(1)}$:

$$C(P^{(1)}, F) = \sum_{i_1=1}^{N-L} \dots \sum_{i_L=L}^{N} [p^{(1)}]_{i_1,\, \text{argmax}(F_1)} \cdot \dots \cdot [p^{(1)}]_{i_L,\, \text{argmax}(F_L)}$$

Here $\text{argmax}(F_j)$ corresponds to the nucleotide identity at the j:th position in the 1-hot-coded motif $F$. We may add/subtract a near-identical cost component on the basis of the sampled 1-hot-patterns $S_k^{(1)}$ to obtain a mean L1-style loss:

$$C(\{S_1^{(1)}, ..., S_K^{(1)}\}, F) = \frac{1}{K} \sum_{k=1}^{K} \sum_{i_1=1}^{N-L} \dots \sum_{i_L=L}^{N} [s_k^{(1)}]_{i_1,\, \text{argmax}(F_1)} \cdot \dots \cdot [s_k^{(1)}]_{i_L,\, \text{argmax}(F_L)}$$

The exact cost configuration, including the fitness loss, the sequence regularization, and loss coefficients, differ for each of the three applications considered in this paper, and we refer the reader to their corresponding sections below for a detailed definition of each cost function.

#### APA Predictor Model (CNN)

We used the deep learning predictor APARENT from (Bogard et. al., 2019) for both the APA isoform and 3' Cleavage design applications. We retrained a modified version of APARENT on the same data set (the 3.5M random MPRA), now as a single network that predicts both isoform and cleavage proportions (instead of two separate networks), and we used all 12 sub libraries of the MPRA for training (in the paper, 3 libraries were held out as independent test data).

**Data**

The APA data set is a synthetic in-vivo MPRA of >3.5 million randomized APA reporters, grown and measured in HEK293 cells. We refer to (Bogard et. al., 2019) for details about the data set. When retraining APARENT, we used 5% of the data for validation and 5% for testing (10% of the MPRA with highest RNA-Seq read count).

**Predictor Architecture**

The predictor is a convolutional neural network consisting of two convolutional layers separated by a max pooling layer, followed by a dense hidden layer. Finally, two separate parallel dense layers output a sigmoid isoform proportion and a softmax cleavage distribution respectively. The exact architecture is given in the table below:

| **Network Layers** | **Parameters** |
|---|---|
| Sequence Input | 1-Hot Encoding, $\{0, 1\}^{205 \times 4}$ |
| Conv(96, 8, Stride=1)<br>ReLU() | Filters = 96<br>Filter Width = 8<br>Stride = 1 |
| MaxPool(pool=2) | Pool Factor = 2 |
| Conv(128, 6, Stride=1)<br>ReLU() | Filters = 6<br>Filter Width = 6<br>Stride = 1 |
| Dense(256)<br>ReLU()<br>Dropout(0.2) | Neurons = 256<br><br>Dropout = 0.2 |
| Dense(1)<br>Sigmoid()<br>Dense(206)<br>Softmax() | Neurons = 1<br><br>Neurons = 206 |

#### APA Isoform Generation

**Generator Architecture**

The generator network receives as input a minibatch of 32 100-dimensional seed vectors of real-valued numbers in range [-1, 1], and during training we uniformly randomly sampled each seed vector component independently. The output of the generator is a minibatch of nucleotide log probability matrices (each 205 nt long). The exact architecture is given in the table below:

| **Network Layers** | **Parameters** |
|---|---|
| Seed | $\mathcal{R}^{100}$ |
| Class | 1-Hot Encoding |
| Concatenate() | Seed & Class vectors |

| | |
|---|---|
| Dense(8064) | Neurons = 8064 |
| Deconv(256, 7, Stride=2)<br>BatchNorm()<br>ReLU() | Filters = 256<br>Filter Width = 7<br>Stride = 2 |
| Deconv(192, 8, Stride=2)<br>BatchNorm()<br>ReLU() | Filters = 192<br>Filter Width = 8<br>Stride = 2 |
| Deconv(128, 7, Stride=2)<br>BatchNorm()<br>ReLU() | Filters = 128<br>Filter Width = 7<br>Stride = 2 |
| Conv(128, 8, Stride=1)<br>BatchNorm()<br>ReLU() | Filters = 128<br>Filter Width = 8<br>Stride = 1 |
| Conv(64, 8, Stride=1)<br>BatchNorm()<br>ReLU() | Filters = 64<br>Filter Width = 8<br>Stride = 1 |
| Conv(4, 8, Stride=1) | Filters = 4<br>Filter Width = 8<br>Stride = 1 |

**Fitness Objective & Training**

The generator weights of the target-isoform DEN are trained for a total of 25,000 updates using the Adam optimizer. We optimize a symmetric KL-divergence between the predicted and target isoform proportions. In RIFR, the objective is defined in terms of $\bar{y}^{(1)}$ as follows:

$$C_{\text{objective}}(\bar{y}^{(1)}) = KL(\bar{y}^{(1)}||t) + KL(t||\bar{y}^{(1)})$$

In SIFR, it is defined as the average symmetric KL-divergence across all K sampled predictions:

$$C_{\text{objective}}(\{y_1^{(1)}, ..., y_K^{(1)}\}) = 1/K \cdot \sum_{k=1}^{K} KL(y_i^{(1)}||t) + KL(t||y_i^{(1)})$$

#### APA Inverse Regression

**Generator Architecture**

The generator network receives as input a minibatch of 32 100-dimensional seed vectors of real-valued numbers in range [-1, 1], which were uniformly sampled during training. The network also receives target isoform log odds values (scalars), which were randomly sampled in the range [-4, 6] during training. The output of the generator is a minibatch of nucleotide log probability matrices (each 205 nt long). The exact architecture is given in the table below:

| Network Layers | Parameters |
|---|---|
| Seed | $\mathcal{R}^{100}$ |
| Target Logit<br>Dense(256)<br>BatchNorm()<br>ReLU()<br>Dense(256)<br>BatchNorm()<br>ReLU() | $\mathcal{R}$<br>Neurons = 8064<br><br><br>Neurons = 8064 |
| Concatenate() | Seed & Logit Encoding vectors |
| Dense(8064) | Neurons = 8064 |
| Deconv(256, 7, Stride=2)<br>BatchNorm()<br>ReLU() | Filters = 256<br>Filter Width = 7<br>Stride = 2 |
| Deconv(192, 8, Stride=2)<br>BatchNorm()<br>ReLU() | Filters = 192<br>Filter Width = 8<br>Stride = 2 |
| Deconv(128, 7, Stride=2)<br>BatchNorm()<br>ReLU() | Filters = 128<br>Filter Width = 7<br>Stride = 2 |
| Conv(128, 8, Stride=1)<br>BatchNorm()<br>ReLU() | Filters = 128<br>Filter Width = 8<br>Stride = 1 |
| Conv(64, 8, Stride=1)<br>BatchNorm()<br>ReLU() | Filters = 64<br>Filter Width = 8<br>Stride = 1 |
| Conv(4, 8, Stride=1) | Filters = 4<br>Filter Width = 8<br>Stride = 1 |

**Fitness Objective & Training**

The generator weights of the inverse-regression DEN are trained for a total of 25,000 updates using the Adam optimizer. We optimize a symmetric KL-divergence between the predicted and target isoform proportions. Note that the target isoform proportion $t$ is a random proportion sampled during trained and is jointly passed as input to the generator. In RIFR, the objective is defined in terms of $\bar{y}^{(1)}$ as follows:

$$C_{\text{objective}}(\bar{y}^{(1)}) = KL(\bar{y}^{(1)}||t) + KL(t||\bar{y}^{(1)})$$

In SIFR, it is defined as the average symmetric KL-divergence across all K sampled predictions:

$$C_{\text{objective}}(\{y_1^{(1)}, ..., y_K^{(1)}\}) = 1/K \cdot \sum_{k=1}^{K} KL(y_i^{(1)}||t) + KL(t||y_i^{(1)})$$

#### 3' Cleavage Generation

**Generator Architecture**

The generator network receives as input a minibatch of 32 100-dimensional seed vectors of real-valued numbers in range [-1, 1], and during training we uniformly randomly sampled each seed vector component independently. The network also receives target cut positions (1-hot-encoding of 9 possible positions), which were uniformly randomly sampled during training. The output of the generator is a minibatch of nucleotide log probability matrices (each 205 nt long). The exact architecture is given in the table below:

| **Network Layers** | **Parameters** |
|---|---|
| Seed | $\mathcal{R}^{100}$ |
| Class | 1-Hot Encoding |
| Concatenate() | Seed & Class vectors |
| Dense(8064) | Neurons = 8064 |
| Concatenate()<br>Deconv(256, 7, Stride=2)<br>BatchNorm()<br>ReLU() | Concatenate Class vector<br>Filters = 256<br>Filter Width = 7<br>Stride = 2 |
| Concatenate()<br>Deconv(192, 8, Stride=2)<br>BatchNorm()<br>ReLU() | Concatenate Class vector<br>Filters = 192<br>Filter Width = 8<br>Stride = 2 |
| Concatenate()<br>Deconv(128, 7, Stride=2) | Concatenate Class vector<br>Filters = 128 |

| | |
|---|---|
| BatchNorm()<br>ReLU() | Filter Width = 7<br>Stride = 2 |
| Concatenate()<br>Conv(128, 8, Stride=1)<br>BatchNorm()<br>ReLU() | Concatenate Class vector<br>Filters = 128<br>Filter Width = 8<br>Stride = 1 |
| Concatenate()<br>Conv(64, 8, Stride=1)<br>BatchNorm()<br>ReLU() | Concatenate Class vector<br>Filters = 64<br>Filter Width = 8<br>Stride = 1 |
| Concatenate()<br>Conv(4, 8, Stride=1) | Concatenate Class vector<br>Filters = 4<br>Filter Width = 8<br>Stride = 1 |

**Fitness Objective & Training**

The generator weights of the target-cleavage DEN are trained for a total of 25,000 updates using the Adam optimizer. We optimize the KL-divergence between the predicted and target cleavage distributions. Note that the target cleavage distribution $\vec{t}$ is the one-hot encoding of a randomly sampled cut position. The sampled cut position is jointly passed as input to the generator. In RIFR, the objective is defined in terms of $\bar{y}^{(1)}$ as follows:

$$C_{\text{objective}}(\bar{y}^{(1)}) = KL(\bar{y}^{(1)}||\vec{t})$$

In SIFR, it is defined as the average KL-divergence across all K sampled predictions:

$$C_{\text{objective}}(\{y_1^{(1)}, ..., y_K^{(1)}\}) = 1/K \cdot \sum_{k=1}^{K} KL(y_i^{(1)}||\vec{t})$$

#### Maximal Polyadenylation Reporter Assay

To evaluate the performance of the DEN-generated pA sequences, we experimentally compared two of the strongest newly generated sequences to two of the strongest gradient ascent-generated sequences from (Bogard et. al., 2019).

**Experiment**

Each of the two newly generated pA sequences were synthesized on plasmid reporters with one of the gradient ascent-sequences, such that they competed for polyadenylation when expressed in cells. Proximal pA signals have a preferential bias of selection than distal pA signals, so to discount this phenomenon we synthesized two orientations of each reporter: (1) The newly generated pA signal proximal and the gradient ascent-signal distal, and (2) The gradient ascent-signal proximal and the new signal distal.

We are interested in estimating the odds ratio of proximal isoform abundance of reporter 1 with respect to reporter 2, as this is equivalent to the fold change in selection preference for the newly generated signals. We estimated this number using a qPCR assay. The plasmids were transfected in HEK293 cells and expressed for 48 hours before RNA extraction. Using two primers per reporter, one priming both possible APA isoforms and one priming only distal isoforms, we used qPCR to read out the corresponding Ct values.

**Ct Difference Lower Bound On Isoform Odds Ratio**

For each pair of reporters, we measure four different qPCR cycle threshold values: (1) $\mathrm{Ct}_1^{all}$, the Ct for both proximal and distal RNA in reporter orientation 1, (2) $\mathrm{Ct}_1^{distal}$, the Ct for both distal RNA only in reporter orientation 1, (3) $\mathrm{Ct}_2^{all}$, the Ct for both proximal and distal RNA in reporter orientation 2, (4) $\mathrm{Ct}_2^{distal}$, the Ct for both distal RNA only in reporter orientation 2. Each cycle threshold value is inversely proportional to the $\log_2$ of the RNA count, and so we can compute the following $\Delta\mathrm{Ct}$ values of log ratios:

$$\Delta\mathrm{Ct}_1 = -(\mathrm{Ct}_1^{\mathrm{all}} - \mathrm{Ct}_1^{\mathrm{distal}}) = \log_2(\frac{p_1 + d_1}{d_1}), \quad \Delta\mathrm{Ct}_2 = -(\mathrm{Ct}_2^{\mathrm{all}} - \mathrm{Ct}_2^{\mathrm{distal}}) = \log_2(\frac{p_2 + d_2}{d_2})$$

Here, $p_1$ and $d_1$ are the proximal and distal RNA count of reporter orientation 1, $p_2$ and $d_2$ are the proximal and distal RNA count of reporter orientation 2, and $\Delta\mathrm{Ct}_1$ and $\Delta\mathrm{Ct}_2$ are the two reporters' respective isoform log ratios. In the rest of this section, we prove that $2^{-\Delta\Delta\mathrm{Ct}} = 2^{-(\Delta\mathrm{Ct}_2 - \Delta\mathrm{Ct}_2)}$ is a lower bound on the isoform fold change, or Odds Ratio, of reporter 1 w.r.t. reporter 2. Using the definitions of $\Delta\mathrm{Ct}_1$ and $\Delta\mathrm{Ct}_2$ above, we have:

$$2^{-\Delta\Delta\mathrm{Ct}} = 2^{-(\Delta\mathrm{Ct}_2 - \Delta\mathrm{Ct}_1)} = \frac{p_1 + d_1}{d_1} / \frac{p_2 + d_2}{d_2}$$

Next, define variables $x_1$ and $x_2$, the proximal isoform proportions of reporter 1 and 2:

$$x_1 = \frac{p_1}{p_1 + d_1}, \quad x_2 = \frac{p_2}{p_2 + d_2}$$

We can rewrite our expression for $2^{-\Delta\Delta\mathrm{Ct}}$ using only these variables:

$$\frac{p_1 + d_1}{d_1} / \frac{p_2 + d_2}{d_2} = \frac{1}{1 - x_1} / \frac{1}{1 - x_2} \quad \frac{p_1 + d_1}{d_1} / \frac{p_2 + d_2}{d_2} = \frac{1}{1 - x_1} / \frac{1}{1 - x_2}$$

Next, we multiply both of the smaller fractions with $x_1$ and $x_2$ on both sides, respectively:

$$= (\frac{1}{x_1}\frac{x_1}{1-x_1})/(\frac{1}{x_2}\frac{x_2}{1-x_2}) = \frac{x_2}{x_1} * (\frac{x_1}{1-x_1}/\frac{x_2}{1-x_2})$$

We can now solve for the Odds Ratio in terms of $2^{-\Delta\Delta \mathrm{Ct}}$. At first glance, it appears we get a rest term $x_1/x_2$ that we cannot solve. However, we know from our experiments that $2^{-\Delta\Delta \mathrm{Ct}} > 1$ and we previously derived that $2^{-\Delta\Delta \mathrm{Ct}} = \frac{1}{1-x_1}/\frac{1}{1-x_2}$. In order for both these equations to be true, then $x_1 > x_2$, and as a consequence $x_1/x_2 > 1$. Hence, our $2^{-\Delta\Delta \mathrm{Ct}}$ measurements are proven to be lower bounds of the isoform odds ratio:

$$\mathrm{Odds\ Ratio}(x_1, x_2) = \frac{x_1}{1-x_1}/\frac{x_2}{1-x_2} = \frac{x_1}{x_2} \cdot 2^{-\Delta\Delta \mathrm{Ct}} >= 2^{-\Delta\Delta \mathrm{Ct}}$$

#### Splicing Predictor Model (CNN)

A deep learning predictor almost identical to the APA predictor was trained on cell-type specific 5' alternative splicing data. The network predicts four splice donor usage proportions, one for each of the cell types HEK, HELA, MCF7 and CHO.

**Data**

We trained the predictor on the random 5' splicing MPRA of (Rosenberg et. al., 2015). The 2015 paper already had data available for HEK293 cells. We replicated the assay according to the protocol described in the original paper, now in cell lines HELA, MCF7 and CHO. For details on the MPRA protocol, we refer to (Rosenberg et. al., 2015). In short, the data consists of approximately 265,000 randomized splicing reporters with alternative 5' splice donors. 25 nt downstream of each splice donor is randomized, for a total of 50 nt of variable sequence. We trained the predictor network on 90% of the data, keeping 5% for validation and 5% for testing (10% of the data with the highest average RNA-Seq read count).

**Predictor Architecture**

The predictor is a convolutional neural network consisting of a subnetwork with two convolutional layers separated by a max pooling layer. This subnetwork is applied on the two separate (1-hot-coded) sequence regions. The two activation maps are concatenated and passed to a dense hidden layer. Next, a dense layer with four sigmoid neurons output the cell-type specific splice donor usage predictions. The architecture is given in the table below:

| **Network Layers** | **Parameters** |
|---|---|
| Sequence Input Region A | 1-Hot Encoding, $\{0,1\}^{35\times4}$ |

| | |
|---|---|
| Sequence Input Region B | 1-Hot Encoding, $\{0,1\}^{35\times4}$ |
| Conv(96, 8, Stride=1)<br>ReLU() | Filters = 96<br>Filter Width = 8<br>Stride = 1 |
| MaxPool(pool=2) | Pool Factor = 2 |
| Conv(128, 6, Stride=1)<br>ReLU() | Filters = 6<br>Filter Width = 6<br>Stride = 1 |
| Concatenate(Region A, Region B) | |
| Dense(256)<br>ReLU()<br>Dropout(0.2) | Neurons = 256<br><br>Dropout = 0.2 |
| Dense(4)<br>Sigmoid() | Neurons = 4 |

#### Splicing Predictor Model (Logistic Regression)

We also trained four logistic regression models, one for each of the cell types HEK, HELA, MCF7 and CHO, to predict the splice donor usage proportions given position-invariant hexamer counts as input features. The same data set that was used to train the splicing neural network was used to train the regression models, including the same training, validation and test splits.

**Predictor Architecture**

We reduce the hexamer count features to differentiable tensor operations by creating a convolutional layer with 4096 orthogonal hexamer detection filters (each filter having a weight of +1 for every encoded nucleotide and an intercept term of -5), and reduce each filter activation map by a sum to obtain a differentiable relaxation of position-invariant hexamer counts extracted from the 1-hot-coded input regions. Each region is separately scored by the convolutional subnet, resulting in a total of 8192 aggregated counts. These counts are weighted according to the cell type-specific logistic regression weights (previously obtained by training on the MPRA), and a final sigmoidal transform becomes the cell type-specific predicted splice donor usage.

| **Network Layers** | **Parameters** |
|---|---|
| Sequence Input Region A | 1-Hot Encoding, $\{0,1\}^{35\times4}$ |

| | |
|---|---|
| Sequence Input Region B | 1-Hot Encoding, $\{0,1\}^{35\times 4}$ |
| Conv(4096, 6, Stride=1)<br>ReLU() | Filters = 4096<br>Filter Width = 6<br>Stride = 1<br>Filter Weight = 6, Bias = -5 |
| Sum() | Over positions |
| Dense(4)<br>Sigmoid() | Neurons = 4 |

#### Differential Splicing Generation

**Generator Architecture**

The generator network followed the architecture that was used for APA isoform generation, except that the final nucleotide log probability matrices produced by the generator were 109 nucleotides long, not 205.

**Fitness Objective & Training**

The generator weights of the max-differential splicing DEN are trained for a total of 25,000 updates using the Adam optimizer. We maximize the difference between the predicted MCF7 and CHO splice donor usages with an absolute difference loss. In RIFR, the objective is defined in terms of $\bar{y}^{(1)}_{\text{CHO}}$ and $\bar{y}^{(1)}_{\text{MCF7}}$ as follows:

$$C_{\text{objective}}(\bar{y}^{(1)}) = 1 - |\bar{y}^{(1)}_{\text{CHO}} - \bar{y}^{(1)}_{\text{MCF7}}|$$

In SIFR, it is defined as the average absolute difference across all K sampled predictions:

$$C_{\text{objective}}(\{y^{(1)}_1, ..., y^{(1)}_K\}) = 1/K \cdot \sum_{k=1}^{K} 1 - |y^{(1)}_{i,\text{CHO}} - y^{(1)}_{\text{i, MCF7}}|$$

### QUANTIFICATION AND STATISTICAL ANALYSIS

We use R^2 as the standard metric for correlation testing throughout the paper. We define R^2 as the square of Pearson's r ($R^2 = r^2$). The exception is when evaluating fitness predictor model performance on each respective application's test set, in which case we define R^2 as the fraction of explained variance ($R^2 = 1 - \mathrm{SSE}/\mathrm{SST}$).

We use two different evaluation metrics when assessing the diversity of a trained DEN generator (i.e. the diversity of the generated sequences). First, we define Duplication Rate $\rho_{\mathrm{dup}}$ as the frequency by which we observe duplicated sequences generated by the model. If we generate $n$ sequences in total and $n_{\mathrm{unique}}$ sequences are unique, we calculate $\rho_{\mathrm{dup}}$ as:

$$\rho_{\mathrm{dup}} = 1 - n_{\mathrm{unique}}/n$$

The second metric we use is entropy. While average single-nucleotide entropy at each position sounds like a reasonable metric, it can sometimes be less expressive than we would like. For example, a trivial generator that continuously shifts a single sequence by one nucleotide at a time could receive a very nucleotide entropy (depending on the sequence). We are more interested in whether the generator is diverse in terms of higher-order motifs. To that end, given a set of sampled sequences, we sum up the total counts of every observed hexamer across the set (regardless of exact position), and obtain a hexamer probability density by normalizing the counts. Next, we compute the Hexamer Entropy in bits as:

$$H_{\mathrm{hex}} = -\sum_{i=1}^{4096} p_i \cdot log_2(p_i)$$

If hexamers where generated entirely uniformly, $H_{\mathrm{hex}}$ would take on its maximum value of 12 bits. If only one hexamer was every generated (e.g. the generator trivially outputs polyA-stretches), the entropy would be 0 bits.

### DATA AND SOFTWARE AVAILABILITY

All code for DEN training and analysis are available on GitHub (https://github.com/johli/genesis). The splicing MPRA data and predictor model are available at https://github.com/johli/splirent.

### KEY RESOURCES TABLE

| **REAGENT or RESOURCE** | **SOURCE** | **IDENTIFIER** |
|---|---|---|
| Deposited Data | | |
| 5' Alternative Splicing MPRA | This study | https://github.com/johli/splirent |
| Software and Algorithms | | |
| DEN Software & Analysis | This study | https://github.com/johli/genesis |
| APARENT Predictor Software | Bogard et. al., 2019 | https://github.com/johli/aparent |
| SPLIRENT Predictor Software | This study | https://github.com/johli/splirent |
