## Supplementary figures and images for "Deep exploration networks for rapid engineering of functional DNA sequences"

### Supplemental Movie 1

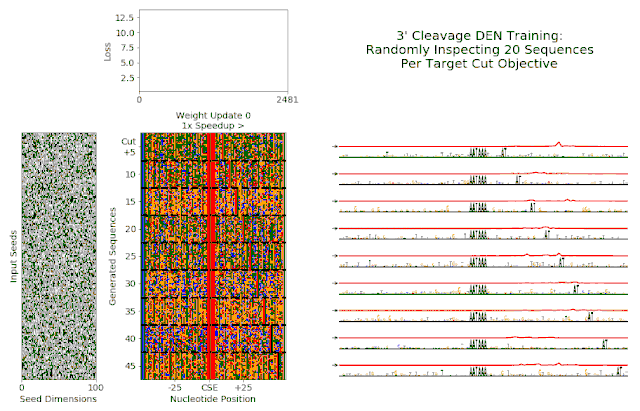

### Supplemental Movie 2

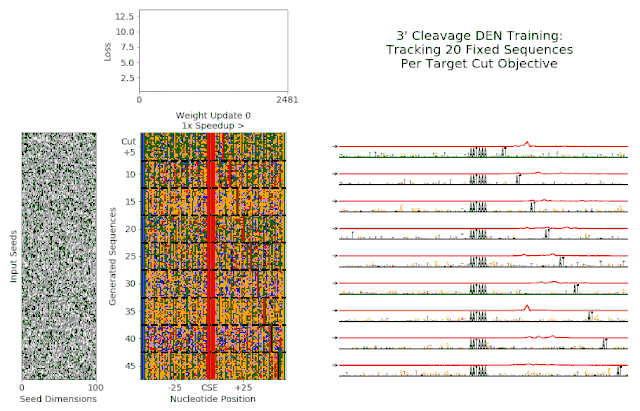
